# Immune landscape of tertiary lymphoid structures in hepatocellular carcinoma (HCC) treated with neoadjuvant immune checkpoint blockade

**DOI:** 10.1101/2023.10.16.562104

**Authors:** Daniel H. Shu, Won Jin Ho, Luciane T. Kagohara, Alexander Girgis, Sarah M. Shin, Ludmila Danilova, Jae W. Lee, Dimitrios N. Sidiropoulos, Sarah Mitchell, Kabeer Munjal, Kathryn Howe, Kayla J. Bendinelli, Hanfei Qi, Guanglan Mo, Janelle Montagne, James M. Leatherman, Tamara Y. Lopez-Vidal, Qingfeng Zhu, Amanda L. Huff, Xuan Yuan, Alexei Hernandez, Erin M. Coyne, Neeha Zaidi, Daniel J. Zabransky, Logan L. Engle, Aleksandra Ogurtsova, Marina Baretti, Daniel Laheru, Jennifer N. Durham, Hao Wang, Robert Anders, Elizabeth M. Jaffee, Elana J. Fertig, Mark Yarchoan

**Author notes:** Corresponding authors: Mark Yarchoan Elana J. Fertig.

## Abstract

Neoadjuvant immunotherapy is thought to produce long-term remissions through induction of antitumor immune responses before removal of the primary tumor. Tertiary lymphoid structures (TLS), germinal center-like structures that can arise within tumors, may contribute to the establishment of immunological memory in this setting, but understanding of their role remains limited. Here, we investigated the contribution of TLS to antitumor immunity in hepatocellular carcinoma (HCC) treated with neoadjuvant immunotherapy. We found that neoadjuvant immunotherapy induced the formation of TLS, which were associated with superior pathologic response, improved relapse free survival, and expansion of the intratumoral T and B cell repertoire. While TLS in viable tumor displayed a highly active mature morphology, in areas of tumor regression we identified an involuted TLS morphology, which was characterized by dispersion of the B cell follicle and persistence of a T cell zone enriched for ongoing antigen presentation and T cell-mature dendritic cell interactions. Involuted TLS showed increased expression of T cell memory markers and expansion of CD8^+^ cytotoxic and tissue resident memory clonotypes. Collectively, these data reveal the circumstances of TLS dissolution and suggest a functional role for late-stage TLS as sites of T cell memory formation after elimination of viable tumor.

**Graphical Abstract:** 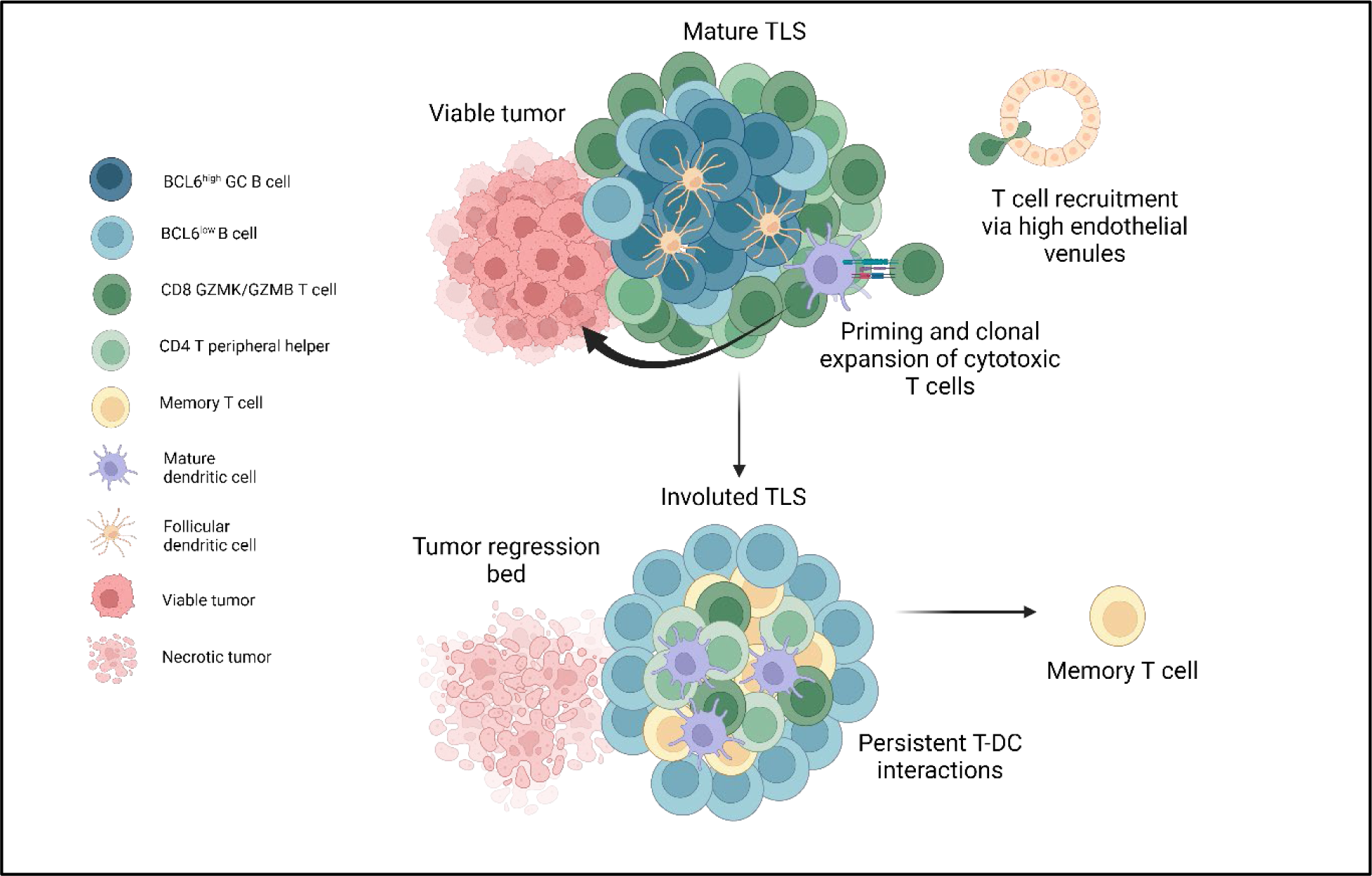

Highlights
1. In patients with hepatocellular carcinoma (HCC), tertiary lymphoid structures (TLS) are induced by neoadjuvant immunotherapy and are associated with favorable clinical outcomes.
2. TLS within the same tumor demonstrate extensive sharing of expanded granzyme K and granzyme B-expressing CD8^+^ T effector memory clonotypes, but the B cell repertoires of individual TLS are almost wholly distinct, consistent with independent germinal center reactions.
3. Within areas of viable tumor, mature TLS are characterized by high expression of CD21 and CD23, BCL6^+^ germinal center B cells, and close interactions between DCLAMP^+^ mature dendritic cells and CXCR5^-^CXCR3^+^ CD4 T peripheral helper cells within a T cell zone adjacent to the B cell follicle.
4. Within areas of tumor regression, an involuted TLS morphology is identified that is notable for dissolution of the B cell germinal center, retention of the T cell zone, and increased T cell memory.

## INTRODUCTION

Immune checkpoint blockade (ICB) therapy has revolutionized the treatment of metastatic solid tumors, offering to a subset of patients the potential for sustained remissions beyond what was previously possible with chemotherapy alone.^1,2^ For patients with early stage, non-metastatic disease, the role for ICB and ideal timing of its administration remains an area of intense clinical investigation. Recent clinical data in patients with melanoma suggest that neoadjuvant immunotherapy, in which ICB is administered prior to curative-intent resection of the primary tumor, may produce superior long-term outcomes compared to immunotherapy given after surgery.^3,4^ Preclinical data suggest that these improved outcomes may be attributable to an elevated and sustained tumor-specific immune response that occurs when immunotherapy is initiated with the primary tumor *in situ*.^5,6^ However, in human subjects it is not known where or by what means the establishment of immunological memory occurs.

Tertiary lymphoid structures (TLS), organized collections of B and T cells that can arise within solid tumors, have been associated with favorable responses to neoadjuvant ICB,^7–13^ and it is hypothesized that TLS play a mechanistic role in promoting effective antitumor immunity. However, understanding of the structure, constituent immune populations, and life cycle of TLS in this treatment setting remains limited by the rarity of neoadjuvant clinical trial specimens, particularly in solid tumor types where the successes of immunotherapy have been modest, and paucity of animal models for TLS in cancer.^14,15^ Thus patient samples from neoadjuvant clinical trials provide a unique opportunity to investigate the contribution of TLS to the development of antitumor immunity.

We previously reported an association between TLS and pathologic response in a phase 1 trial of patients with locally advanced HCC who received neoadjuvant nivolumab and cabozantinib.^8^ Here, we evaluated the clinical and immunological characteristics of TLS in an expanded cohort of patients with locally advanced hepatocellular carcinoma (HCC) treated with neoadjuvant ICB. We find evidence that neoadjuvant ICB induces the formation of intratumoral TLS, and that high TLS density following neoadjuvant therapy is associated with superior pathologic response to treatment and relapse-free survival. Using a multiomics approach employing imaging mass cytometry, bulk TCR and BCR sequencing of microdissected TLS, and paired single cell RNA and TCR sequencing, we identify key differences in the spatial and immunological landscape of TLS in areas of viable and nonviable tumor that suggest that the contribution of TLS to antitumor immunity in tumors treated with neoadjuvant immunotherapy varies significantly according to morphological stage and circumstance.

## RESULTS

### Neoadjuvant ICB in HCC induces intratumoral TLS

To determine the clinical significance of TLS in patients with HCC treated with neoadjuvant ICB, we identified patients from the Johns Hopkins Liver Cancer Biorepository who had undergone surgical resection of their primary tumor after receiving neoadjuvant ICB-based therapy for locally advanced HCC. In total, 19 patients were identified who received treatment between October 2019 and January 2022 (**Extended Data Table 1**). 11/19 (57.9%) were male, 13/19 (68.4%) had tumors with moderately differentiated histology, and 11/19 (57.9%) had a history of viral hepatitis. No patients had active viral hepatitis at the time of surgery. 14/19 (73.6%) received anti-PD-1 plus an oral tyrosine kinase inhibitor, 3/19 (15.8%) received anti-PD-1 monotherapy, 1/19 (5.2%) received combination anti-PD-1 and anti-CTLA-4 monoclonal antibody, and 1/19 (5.2%) received combination anti-PD-1/anti-CTLA-4 and oral TKI prior to resection of the primary tumor.

Since TLS are known to occur in treatment naïve HCC,^16^ we first attempted to determine if TLS were present in the tumors of patients prior to receiving neoadjuvant ICB. 7/19 (36.8%) of patients had undergone pre-treatment fine needle biopsies prior to initiation of neoadjuvant therapy and no intratumoral TLS were identified in these specimens. Given the limited assessment of the tumor microenvironment provided by fine needle biopsy, we next identified a second cohort of HCC patients treated at our institution who had undergone surgical resection without receiving prior systemic therapy, which would serve as a control cohort. 17 patients were identified who had received upfront surgical resection for HCC between 2017 and 2022, from which 3 patients were excluded due to small tumor volume, poor tissue quality, or HCC etiology not represented by the treatment cohort. The 14 remaining patients (**Extended Data Table 2**), were similar to the neoadjuvant treatment cohort by age, sex, histologic grade, and etiology (**Table 1**).

**Table 1.** Characteristics of the Patients, According to Treatment Status.

|  | Neoadjuvant<br>(N=19) | Untreated<br>(N=14) | Total<br>(N=33) |
| --- | --- | --- | --- |
| **Age at surgery — yr** |  |  |  |
| Mean (SD) | 64±10 | 66±9.0 | 65±9.6 |
| Median (range) | 65 (41-79) | 66 (49-84) | 65 (41-84) |
| **Sex — no. (%)** |  |  |  |
| Male | 11 (58) | 11 (79) | 22 (67) |
| Female | 8 (42) | 3 (21) | 11 (33) |
| **Histologic grade — no. (%)** |  |  |  |
| Poorly differentiated | 3 (16) | 1 (7) | 4 (12) |
| Moderately differentiated | 13 (68) | 8 (57) | 21 (64) |
| Well differentiated | 3 (16) | 5 (36) | 8 (24) |
| **Etiology — no. (%)** |  |  |  |
| HBV | 3 (16) | 1 (7) | 4 (12) |
| HCV | 7 (37) | 6 (43) | 13 (39) |
| HBV/HCV | 1 (5) | 1 (7) | 2 (6) |
| NASH | 2 (11) | 3 (21) | 5 (15) |
| ETOH | 1 (5) | 1 (7) | 2 (6) |
| Unknown | 5 (26) | 2 (14) | 7 (21) |
| **Neoadjuvant Treatment — no. (%)** |  |  |  |
| anti-PD1 + TKI | 14 (74) | 0 (0) | 14 (42) |
| anti-PD1 | 3 (16) | 0 (0) | 3 (9) |
| anti-PD1 + anti-CTLA4 | 1 (5) | 0 (0) | 1 (3) |
| anti-PD1 + anti-CTLA4 + TKI | 1 (5) | 0 (0) | 1 (3) |
| None | 0 (0) | 14 (100) | 14 (42) |

Evaluation of TLS density in the two cohorts was performed by CD20 staining of resected FFPE tumor (**Fig. 1a**). TLS, which we defined as CD20^+^ lymphoid aggregates with diameter greater than 150 μm, were classified as either peritumoral or intratumoral according to their location relative to the interface between tumor and normal adjacent parenchyma (**Fig. 1b**). TLS were observed in 7/14 (50%) of untreated tumors and 12/19 (63.2%) treated tumors. No significant difference was identified in total TLS density (0.08±0.09 TLS/mm^2^ versus 0.05±0.10 TLS/mm^2^, P = 0.42) or peritumoral TLS density (0.03±0.05 TLS/mm^2^ versus 0.04±0.1 TLS/mm^2^, P = 0.73) (**Extended Fig. 1a**), but intratumoral TLS density was significantly increased in treated patients compared to untreated controls (0.05±0.08 TLS/mm^2^ versus 0.01±0.02 TLS/mm^2^, P = 0.05) (**Fig. 1c**). In untreated tumors, the majority of TLS were peritumoral, whereas in neoadjuvant treated tumors the majority were intratumoral (**Extended Data Fig 1b-c**). Taken together, these data suggest that neoadjuvant ICB induces the formation of intratumoral TLS.

**Fig. 1.**
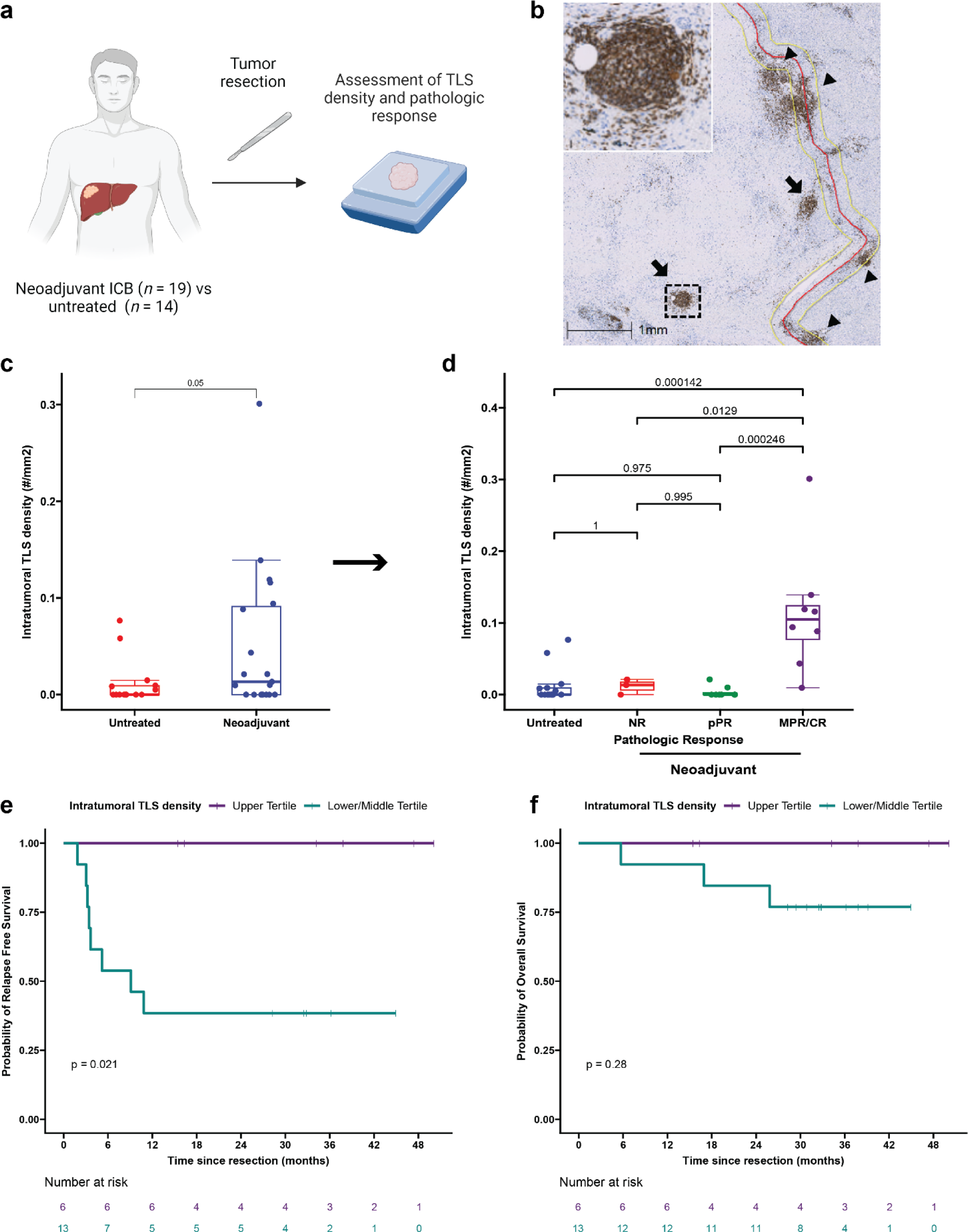
Neoadjuvant ICB induces intratumoral TLS, which are associated with superior pathologic response and relapse free survival. **a**, Workflow for TLS density analysis. **b**, Representative images of formalin fixed paraffin embedded (FFPE) HCC tumors stained with anti-CD20 antibody. Annotations indicate boundary between tumor/tumor regression bed and adjacent normal parenchyma (red), extension of boundary by 200 μm (yellow), intratumoral TLS (arrow), and peritumoral TLS (arrow head). Inset shows representative TLS at high magnification. Scale bar, 1mm. **c**, Box-and-whisker plots showing intratumoral TLS density in patients with locally advanced HCC treated with neoadjuvant ICB (*n* = 19) and untreated controls (*n* = 14). **d**, Boxplot-and-whisker plots showing intratumoral TLS density in untreated (*n* = 14) and neoadjuvant treated tumors, divided according to pathologic response (*n* = 19). For each box-and-whisker plot, the horizontal bar indicates the median, the upper and lower limits of the boxes the interquartile range, and the ends of the whiskers 1.5 times the interquartile range. **e-f**, Kaplan-Meier curves showing relapse free survival (**e**) and overall survival (**f**) for patients with HCC in the highest tertile (purple) compared to the middle and lowest tertiles (green) of intratumoral TLS density after neoadjuvant ICB. Statistical significance was determined by two-tailed t-test (c), one-way ANOVA followed by Tukey’s honest significant difference (HSD) test (**d**), and log-rank test (**e** and **f**).

### High intratumoral TLS density after neoadjuvant ICB is associated with superior pathologic response and disease-free survival

We next set out to determine if there were an association between high TLS density after neoadjuvant ICB and three clinically meaningful endpoints: pathologic response to treatment, relapse free survival, and overall survival. Tumors treated with neoadjuvant ICB were reviewed and assigned to a pathologic response category (non-response [NR], partial pathologic response [pPR], or major or complete pathologic response [MPR/CR]) according to percent residual viable tumor at the time of surgery.^17^ 8/19 (42.1%) patients had a major or complete pathologic response, of which 2 had CR and 6 had MPR; 8/19 (42.1%) had a partial pathologic response (pPR); and 3/19 (15.8%) had non-response (NR). Intratumoral TLS density was significantly increased in tumors with MPR/CR compared to tumors with pPR (P = 0.000246), NR (P = 0.0129), or untreated patients (P = 0.000142) by Tukey’s HSD test. In addition, total TLS density was also increased in tumors with MPR/CR compared to tumors with pPR (P = 0.00144), NR (P = 0.02), and untreated tumors (P = 0.00694) (**Fig. 1d** and **Extended Data Fig. 1d**). No significant difference was observed in peritumoral TLS density across pathologic response groups or untreated tumors (**Extended Data Fig. 1e**). Additional pathologic assessment was also performed according to the Immune Related Pathologic Response Criteria (irPRC), a set of categorical histopathologic criteria developed for standardized pathologic assessment of the regression bed of neoadjuvant immunotherapy treated solid tumors.^17^ Using these criteria, we also observed a significant association between the presence of intratumoral TLS and MPR/CR (P = 0.02), while no significant association was detected between peritumoral TLS and MPR/CR (P = 0.38) (**Extended Data Table 3**). Thus, both a quantitative assessment of TLS density and a categorical evaluation of individual pathologic features suggested that intratumoral TLS density may be most correlated with response to treatment.

We next examined relapse free survival and overall survival in the treated cohort, excluding the untreated cohort from analysis lack of follow up data for the majority of the cohort. Significantly longer relapse free survival after surgery was observed in treated patients in the upper tertile of intratumoral TLS density compared to patients in the middle and lower tertiles (P = 0.021) (**Fig. 1e**). At a median follow up of 38 months for patients in the upper tertile of intratumoral TLS density and 32 months for patients in the middle and lower tertiles, median RFS was not reached in the upper tertile and 9.1 months in the middle and lower group. RFS at 30 months was 100% and 38.5% (95% CI, 19.3% to 76.5%), respectively. No significant difference in overall survival (OS) was observed between the two groups (P = 0.24) (**Fig. 1f**), but at 30 months OS was 100% in the upper tertile and 76.9% (95% CI, 57.1% to 100%) in the middle and lower tertiles. In addition, we observed a trend toward improved RFS for patients in the upper tertile of total TLS density compared to the middle and lower tertiles (P = 0.13) (**Extended Data Fig. 2a**). OS was not significantly different (P = 0.28) (**Extended Data Fig. 2b**), but no deaths were observed in the upper tertile of total TLS density while three deaths were observed in the middle and lower tertiles. With respect to peritumoral TLS density, no difference was observed in relapse free survival (P = 0.56) or overall survival (p = 0.23) when comparing the upper tertile to the middle and lower tertiles (**Extended Data Fig. 2c-d**). Notably, in this cohort MPR/CR, which was closely associated with high TLS density, was also associated with superior RFS (P = 0.025) (**Extended Data Fig. 2e**). No significant difference was observed in OS (P = 0.16) (**Extended Data Fig. 2f**), but no deaths were observed in the MPR/CR group while three deaths were observed in the pPR/NR group. In addition, we also evaluated outcomes according to sex and previous viral HBV or HCV infection and identified no significant differences in RFS or OS (**Extended Data Fig. 2g-l**). Finally, to compare the different clinical covariates, we used the Bayesian information criterion^18^ to quantify the strength of each parameter in predicting relapse free survival or death after neoadjuvant ICB and surgical resection. The strongest predictors of relapse free survival by BIC analysis were intratumoral TLS density and pathologic response (**Extended Data Table 4**).

### High TLS density after neoadjuvant ICB is associated with increased T and B cell activation and an expanded intratumoral T and B cell repertoire

To identify differences in gene expression between tumors with high and low TLS density in this treatment context, we performed bulk RNA sequencing from FFPE surgical resection specimens. Tissue sections were collected from 14 tumors in the neoadjuvant treatment group, of which 2 samples were excluded after quality control. The resultant 12 samples were designated as TLS high (*n* = 5) or TLS low (*n* = 7) according to total TLS density relative to the mean total TLS density of the treatment group. Here, total TLS density was used rather than intratumoral or peritumoral TLS density since bulk sequencing of FFPE tissue blocks did not have spatial resolution to account for these differences. By principal component analysis, the 5 TLS high tumors and 1 TLS low tumor clustered separately from the remaining TLS low tumors (**Fig. 2a**). Differential expression analysis using the R package DESeq2 identified 814 differentially expressed genes (DEG), defined as having fold change in the TLS high group greater than 2 times that of the TLS low group and false discovery rate less than 0.05 (**Fig. 2b-c** and **Extended Data Table 5**).

**Fig. 2.**
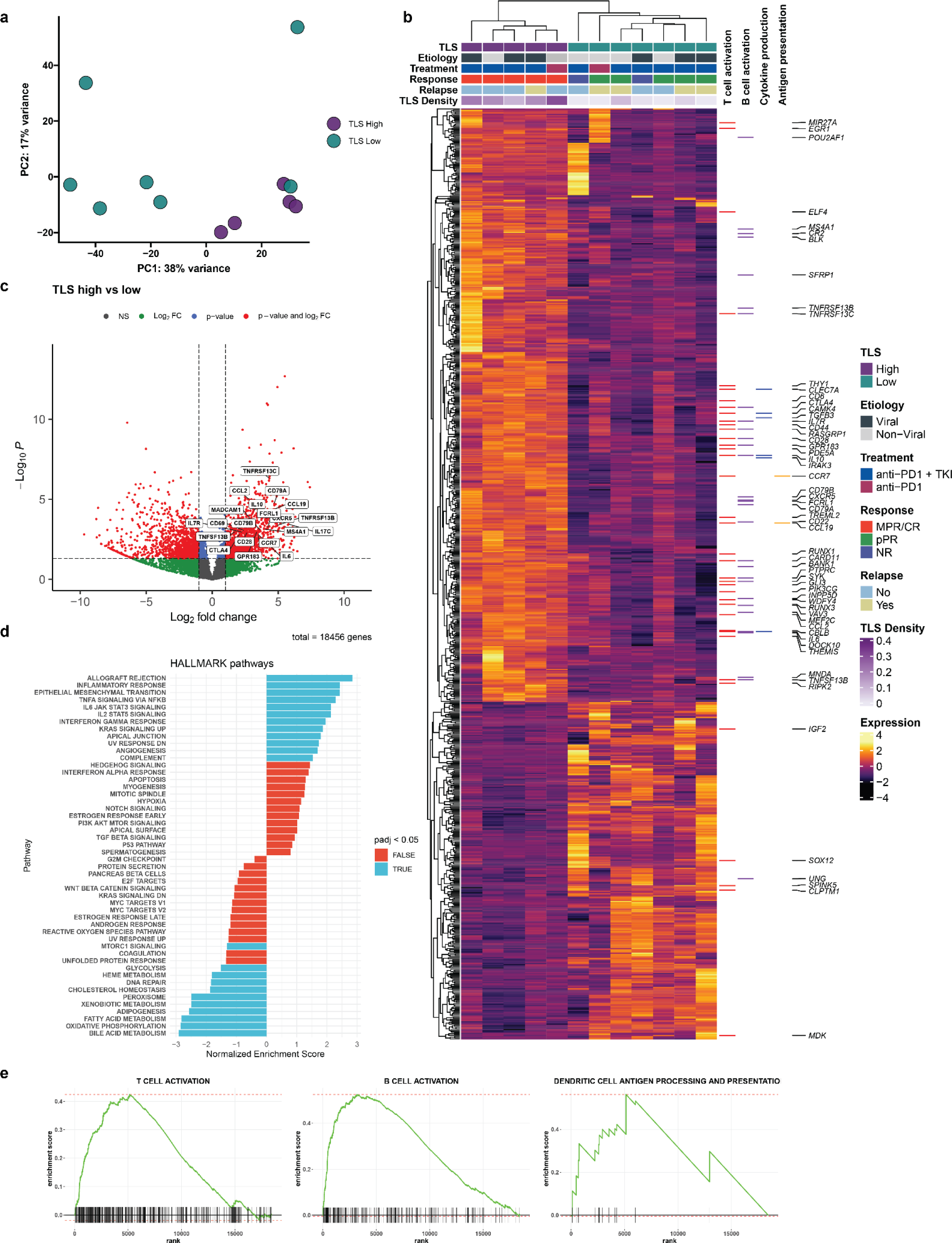
High TLS density is associated with increased T and B cell activation in HCC treated with neoadjuvant ICB. **a**, Principle component analysis of bulk RNA-sequencing of resected HCC tumors treated with neoadjuvant ICB (*n* = 12), divided according to TLS density relative to mean density of the neoadjuvant treatment group. **b,** Heatmap showing differentially expressed genes (DEG) with a log2 fold change > 1 and P < 0.05 between tumors with high (*n* = 5) and low (*n* = 7) TLS density. Annotation rows indicate TLS group, HCC etiology, treatment, response, relapse, and TLS density. Annotation columns at right identify DEG belonging to Gene Oncology Biological Pathways gene sets for T cell activation, B cell activation, Cytokine production, and Dendritic Cell Antigen Processing and Presentation. **c,** Volcano plot showing differentially expressed genes between tumors with high and low TLS density. Vertical dotted lines represents log2 fold change greater than or less than 1. Horizontal dotted line indicates adjusted P value of 0.05. 4 outlier genes are excluded from the plot for the purposes of visualization. **d,** Gene set enrichment analysis showing differentially enriched gene sets from the HALLMARK database between tumors with high and low TLS density. **e,** Barcode plots showing enrichment scores for the Gene Ontology Biological Pathways gene sets for T cell activation, B cell activation, and Dendritic Cell Antigen Processing and Presentation.

Compared to TLS low tumors, TLS high tumors demonstrated significant overexpression of multiple genes belonging to the Gene Ontology Biological Pathways gene sets for T and B cell activation, cytokine production, and antigen presentation, including *CTLA4*, *IL7R*, *IL6*, the B cell activating factor BAFF(*TNFSF13B*) and its receptors BAFF-R (*TNFRSF13C*) and TACI (*TNFRSF13B*), and the T cell-derived cytokine *IL17C*. TLS high tumors displayed significantly greater expression of *CCL19*, a chemokine involved in T-cell and B-cell migration to secondary lymphoid organs, and *CXCR5*, the receptor for the B-cell chemoattractant *CXCL13*. TLS high tumors also demonstrated increased expression of multiple B-cell related genes such as the B cell antigen CD79 (*CD79A* and *CD79B*), CD20 (*MS4A1*), and Fc Receptor Like A protein (*FCRLA*) which is highly expressed in germinal center B cells.^19^ In addition, we found increased expression of immunoregulatory genes including *IL10*, *IL17REL*, and the integrin αvβ8-mediated *ITGB8*, which mediates TGF-beta-1 activation on the surface of regulatory T cells.^20,21^ We also identified significantly increased expression of the gene encoding the germinal center regulatory protein EBI2 (*GPR183*), *DOCK10*, which regulates CD23 expression and sustains B-cell lymphopoiesis in secondary lymphoid tissue,^22^ and *WDFY4*, a mediator of dendritic cell cross presentation.^23^

Gene set enrichment analysis for human gene sets in the MSigDB collections further identified significant enrichment in TLS high tumors of pathways associated with increased adaptive immune response, including Hallmark pathways for allograft rejection and inflammatory response, and multiple pathways related to T and B cell receptor activation (**Fig. 2d-e** and **Extended Data Table 6**). Consistent with these findings, TLS high tumors also displayed increased expression of the 12-chemokine gene signature which has previously been found in association with TLS formation in multiple solid tumor types (**Extended Data Fig. 3a**).^24^ Taken together, these bulk gene expression data demonstrate that tumors with high TLS density display significantly higher levels of T and B cell activation compared to TLS low tumors.

To determine if TLS density was associated with differences in the adaptive immune repertoire, we used the Personalis ImmunoID NeXT platform to extract immunoglobulin heavy chain (IGH), TCRβ, and TCRα repertoire data from bulk RNA sequencing data. Statistical power was limited by the small sample size, but in tumors with high TLS density there were a significant increase in total number of immunoglobulin heavy chain (IGH) clones (P = 0.02), unique clonotypes (P = 0.029), and repertoire diversity (P = 0.043) by Wilcoxon rank sum test (**Extended Data Fig. 4a- c**). In addition, we identified a trend toward increased median number of total clones, unique clonotypes, repertoire diversity in the TCRα (P *=* 0.29, 0.18, and 0.18, respectively, by Wilcoxon rank sum test) (**Extended Data Fig. 4d-f**) and TCRβ repertoires (P *=* 0.22, 0.095, and 0.095, respectively, by Wilcoxon rank sum test) (**Extended Data Fig. 4g-i**). Overall, these findings suggest that high TLS density is associated with an expansion of the B and T cell repertoire in HCC treated with neoadjuvant immunotherapy.

### In areas of tumor regression, an involuted TLS morphology is found that displays dissolution of the B cell germinal center, retention of the T cell zone, and increased expression of T cell memory markers

Based on these data, we hypothesized that a distinctive immunological process may occur in tumors with high intratumoral TLS density and major or complete pathologic response that contributes to long-term disease-free survival. To evaluate this hypothesis, we performed histologic examination of tumors with both viable tumor and extensive tumor regression beds. In viable tumor, the predominant phenotype observed was the canonical ‘mature’ stage of TLS characterized by a CD20^+^ B cell germinal center surrounded by CD4^+^ and CD8^+^ T cells.^25^ TLS of this morphology showed characteristically high expression of the follicular dendritic cell marker CD21 and the proliferation marker Ki67. In contrast, in areas of tumor regression bed we observed an ‘involuted’ TLS morphology characterized by CD20^+^ B cells in a halo-like ring surrounding a central core of CD4^+^ and CD8^+^ T cells. CD21 and Ki67 expression were low to absent (**Fig. 3a**). To confirm that this involuted morphology was not an artifact of sectioning, we performed serial sectioning and anti-CD20 staining of FFPE tissue sections and confirmed the absence of a dense B cell core as is seen in mature TLS (**Extended Data Fig. 5a**). These involuted TLS were highly associated with tumors with complete pathologic response, and in several tumors were found in series (**Extended Data Fig 5b**), suggesting a shared lymphatic supply. No TLS of this morphology were detected in untreated tumors.

**Fig. 3.**
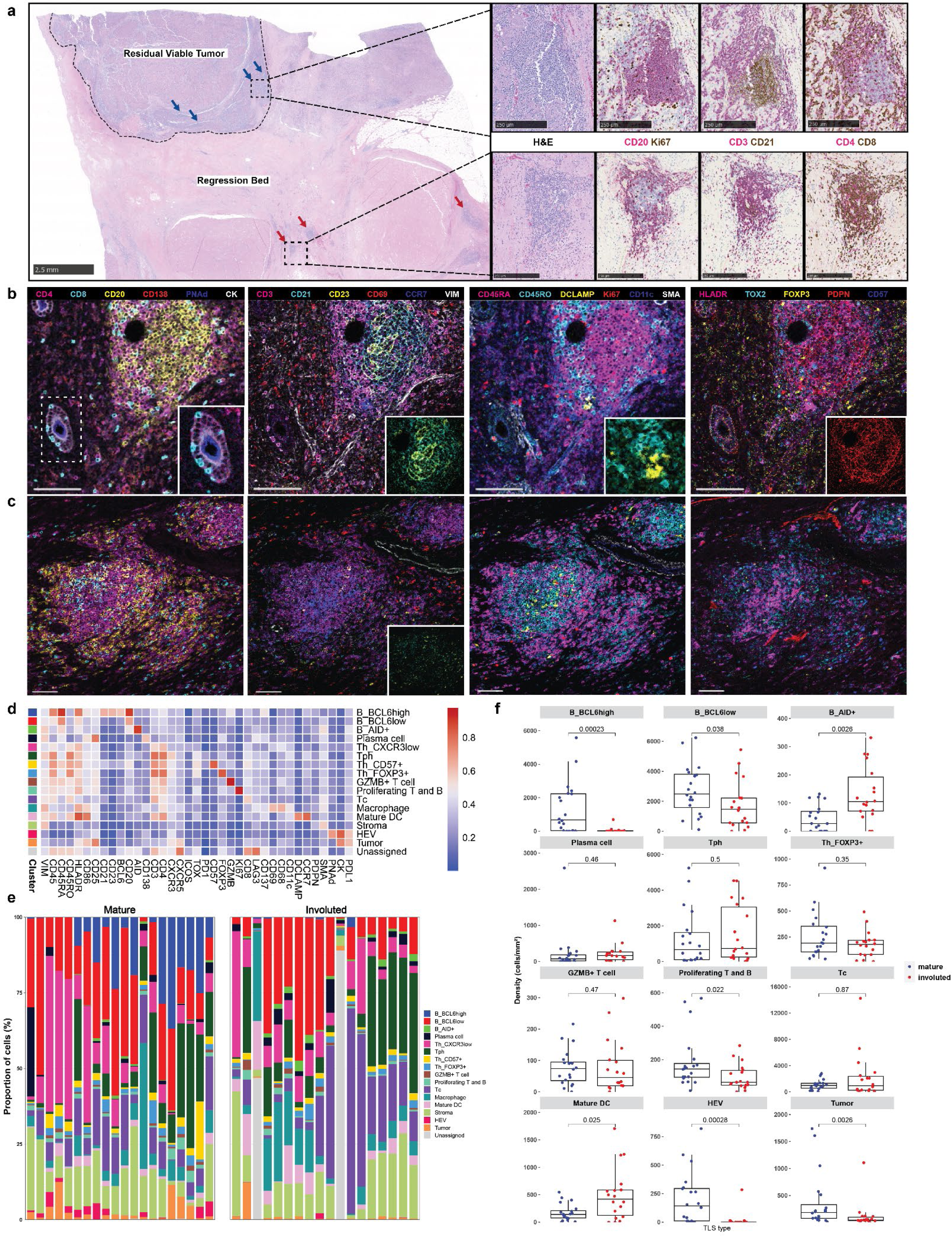
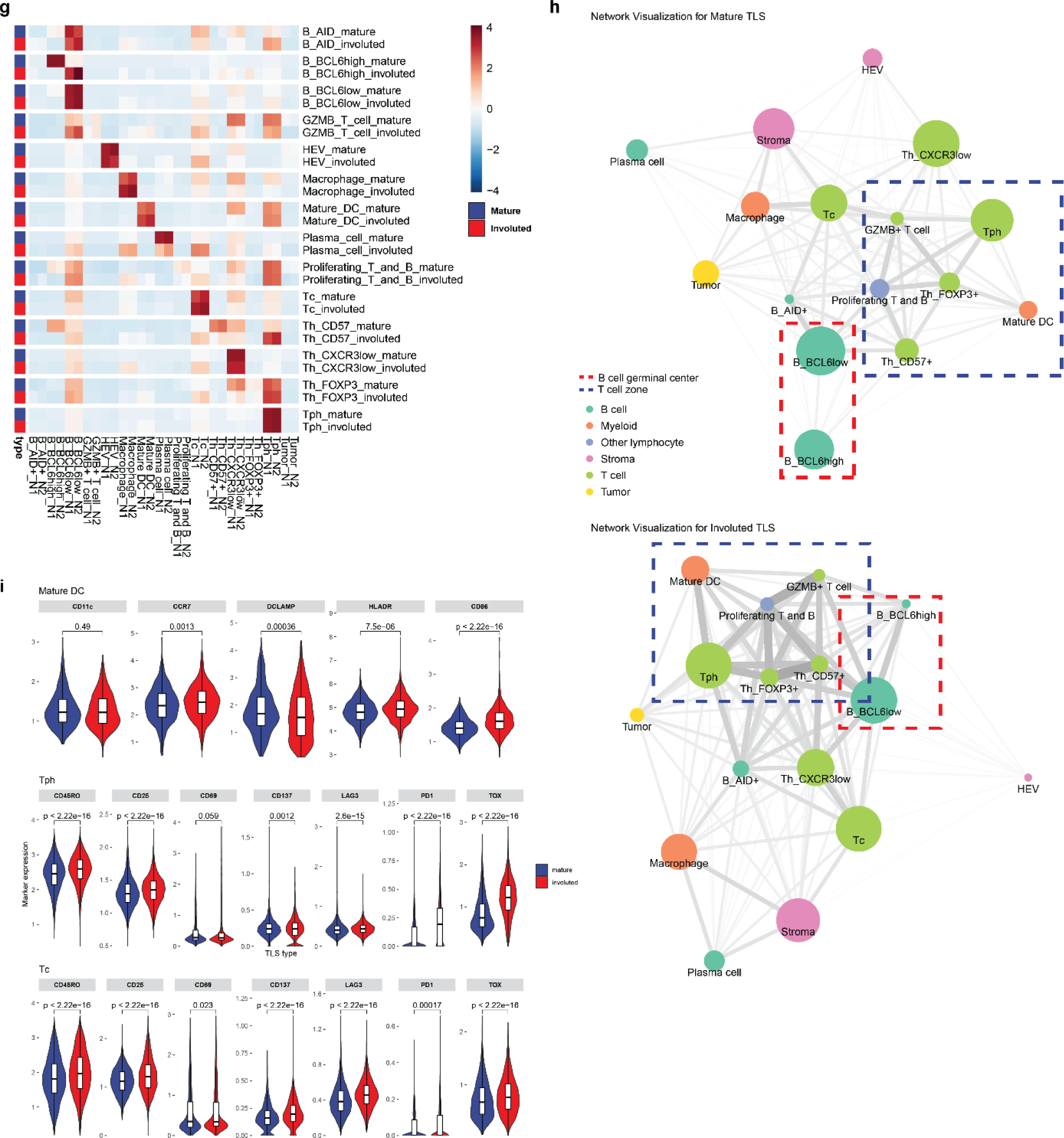
Identification of divergent TLS morphologies and cellular spatial relationships in viable tumor and tumor regression bed. **a**, Representative formalin-fixed, paraffin embedded (FFPE) neoadjuvant ICB-treated tumor stained with hematoxylin and eosin (H&E) showing divergent TLS morphologies (“mature” and “involuted”) in viable residual viable tumor and regression bed. Dotted line shows boundary between residual viable tumor and regression bed. Blue arrows indicate mature TLS and red arrows indicate involuted TLS. Scale bar, 2.5mm. Higher magnification images of representative mature and involuted TLSare shown on right with serial sections stained with dual immunohistochemistry staining for CD20 (magenta) and Ki67 (brown), CD3 (magenta) and CD21 (brown), and CD4 (magenta) and CD8 (brown). Scale bars, 250 μm. **b-c,** Representative images of mature (**b**) and involuted (**c**) TLS obtained by imaging mass cytometry. Insets show higher magnification images of CD8^+^ T cells trafficking through high endothelial venules (**b, far left**), an extensive CD21^+^CD23^+^ follicular dendritic cell network in the mature morphology (**b**, **middle left**) compared to scant CD21^+^ and CD23^+^ in the involuted morphology (**c**, **middle left**), close interactions between T cells and DCLAMP^+^ mature dendritic cells in the T cell zone adjacent to the germinal center (**b, middle right**), and high podoplanin expression in the germinal center of the mature TLS (**b**, **far right**). Scale bars, 100 μm. **d,** Heatmap showing average IMC marker expression in annotated cell clusters identified from 90,344 single cells from 38 TLS (*n =* 20 mature, *n* = 18 involuted). **e,** Composition of mature and involuted TLS regions by cell type as a percentage of total cells per TLS. **f**, Box-and-whisker plots showing cell cluster density in mature versus involuted TLS. For each box-and-whisker plot, the horizontal bar indicates the median, the upper and lower limits of the boxes the interquartile range, and the ends of the whiskers 1.5 times the interquartile range. **g,** Nearest neighbor analysis with rows indicating individual clusters in mature and involuted TLS and columns corresopnding to first and second most common neighbors. **h,** Network analysis for cell clusters in mature and involuted TLS. Node size corresponds to the proportion of total cells for each TLS type occupied by each cluster. Edge length represents the shortest distance between cell clusters and thickness corresponds to the number of measurements for each TLS type. **i**, Box and violin plots showing expression of mature dendritic cell markers (CD11c, CCR7, DCLAMP, HLADR, and CD86) in the mature DC cluster and markers of T cell activation and exhaustion (CD45RO, CD25, CD69, CD137, LAG3, PD1, and TOX) in the T peripheral helper (Tph) and cytotoxic T cell (Tc) clusters, by TLS morphology. Statistical significance was determined by pairwise two sample Wilcoxon test (**f** and **g**).

Given the location of the latter morphology within areas of nonviable tumor and the dispersed appearance of B cells in these lymphoid aggregates, we hypothesized that this morphology may represent TLS undergoing shutdown of the germinal center.^26,27^ To characterize the features of this stage of TLS, we developed a 38-marker imaging mass cytometry antibody panel. Markers included in this panel were selected to identify different T cell subsets (CD3, CD4, CD8, FOXP3, CXCR3, CXCR5, ICOS), B cells subsets (CD20, BCL6, AID, CD138), follicular dendritic cells (CD21, CD23), dendritic cells (CD11c, DC-LAMP, CCR7), high endothelial venules (PNAd), macrophages (CD68), fibroblasts (Podoplanin [PDPN], αSMA), and tumor (CK). We included markers for T cell activation and exhaustion (CD25, CD69, CD137, PD-1, LAG3, TOX), co-stimulatory or antigen presenting molecules (CD86, HLA-DR), and markers of cell proliferation (Ki67) (**Extended Data Tables 7 and 8**). FFPE sections were obtained from the tumors of 9 patients treated with neoadjuvant ICB, in 8 of which the involuted morphology was identified, and, after whole-slide staining, 31 regions of interest (ROI) were captured by laser ablation (**Extended Data Fig. 6a**) from which 38 TLS areas (*n* = 20 mature and 18 involuted) were identified.

Consistent with previously published data, imaging mass cytometry of mature TLS (**Fig. 3b**) demonstrated dense B cell follicle-like structures surrounded by peripherally located CD4^+^ and CD8^+^ T cells with associated high endothelial venules (HEV) with a cuboidal morphology.^28,29^ In multiple HEV, we observed CD8^+^ T cells in transit through these structures (**Fig. 3b, far left inset**). Mature TLS were also notable for an extensive CD21 and CD23 follicular dendritic cell network (**3b, middle left inset**), a distinct T cell zone with densely concentrated DCLAMP^+^ mature dendritic cells in close contact with T cells (**3b, middle right inset**), and a dense PDPN^+^ stromal network, similar to the fibroblastic reticular cell networks seen in secondary lymphoid organs^15,30^ (**3b, far right inset**). Involuted TLS (**Fig. 3c**) showed no detectable HEV with a cuboidal morphology, scattered CD21 and CD23 expression, and diminished PDPN expression, consistent with attenuation of the TLS structure. Notably, the center of involuted TLS demonstrated apparent persistence of the T cell zone with co-location of DCLAMP^+^ mature dendritic cells and CD4^+^ and CD8^+^ T cells.

Further quantitative analysis of these structures supported these initial observations. After cell segmentation, we identified 61,371 single cells which were assigned to 16 distinct cell clusters (**Fig. 3d-f** and **Extended Data Fig. 6b-c**). In mature TLS we observed significantly higher density of a BCL6^high^ population of B cells (B_BCL6^high^), which was consistent with a germinal center B cell population (P = 0.00023). In mature TLS, this cluster localized to the center of the B cell germinal center in close proximity to CD21^+^CD23^+^ follicular dendritic cells and demonstrated high expression of HLADR, a marker of antigen presentation, and CD86, a B cell activation marker, consistent with an activate B cell population. A second B cell cluster was identified on the periphery of the B cell follicle which displayed lower expression of BCL6 (B_BCL6^low^) and decreased expression of HLADR and CD86. This cluster was also found in significantly higher density in mature TLS (P = 0.038). In contrast, in involuted TLS we observed significantly increased density of a third B cell cluster (B_AID+) (P = 0.0026), which was characterized by high expression of activation-induced cytidine deaminase (AID), the B cell enzyme which drives somatic hypermutation and class switch recombination. AID is induced by BCR cross-linking and has a half-life of 2.5 hours in the nucleus and 18-20 hours in its cytoplasmic form,^31,32^ thus this population may correspond to B cells undergoing somatic hypermutation or memory B cells which had recently undergone immunoglobulin class switching, the latter of which we felt to be more likely given the context. No difference was observed in plasma cell densities between the two morphologies.

In the T cell compartment, we identified a single cytotoxic CD8^+^ T cell population (Tc) and two major CD4 T helper populations, a CD4^+^CXCR5^-^CXCR3^+^ T peripheral helper (Tph) cluster, which was located around the peripheral of the B cell germinal center in mature TLS and at the center of involuted TLS, and a CD4^+^CXCR3^-^ T helper (Th_CXCR3^low^) clusters. In location and marker expression, Tph in these data were consistent with CD4^+^ Tph that have been identified in patients with autoimmune disease, where they are thought to play a T follicular helper (Tfh)-like role in promoting pathogenic B cell responses in non-lymphoid tissue.^33–35^ In contrast to changes observed in the B cell compartment, no significant difference was observed in density of CD4^+^ T cell clusters or the cytotoxic Tc cluster.

Clustering analysis also identified a CD4^+^CD57^+^ cluster (Th_CD57^+^) within the germinal center of mature TLS, which may provide help to B cells and induce class switch recombination,^36–38^ a cluster of CD4^+^FOXP3^+^ regulatory T cells (Th_FOXP3^+^), a cluster of cytotoxic T cells (Tc); CD4^+^ and CD8^+^ T cells defined by high expression of granzyme B (GZMB^+^ T cell); proliferating T and B cells defined by high expression of Ki67 (Proliferating T and B); a macrophage cluster with high expression of CD68; a mature dendritic cell cluster defined by the presence of high expression of DCLAMP and CCR7^39^; a high endothelial venule (HEV) cluster defined by expression of the protein peripheral node addressin (PNAd); and a tumor cluster with high expression of Cytokeratin (CK) and PDL1. In mature TLS compared to involuted TLS, there was significantly higher density of proliferating T and B cells (P = 0.022), HEV (P = 0.00028), and tumor (P = 0.0026). On the other hand, density of mature dendritic cells was increased in involuted TLS (P = 0.025).

To further evaluate the spatial relationships between different cell types in the mature and involuted morphologies, we performed nearest neighbor analysis of the top 2 most frequent cell neighbors for each cell cluster (**Fig. 3g**). As in previous the above analyses, neighborhood analysis showed that the primary differences in spatial relationships between the two morphologies occurred in B cell clusters. In particular, in the mature morphology BCL6^high^ germinal center B cell cluster were first and second nearest neighbors for themselves, consistent with a highly concentrated germinal center. In contrast, in the involuted TLS morphology the most common first neighbor of this cluster was the BCL6^low^ cluster, consistent with a more dispersed germinal center in the involuted TLS. On the other hand, Tph were the most common non-self neighbors for GZMB T cells, mature dendritic cells, proliferating T cells, and FOXP3^+^ Tregs in both mature and involuted TLS, suggesting that the spatial relationships of these clusters was preserved across the two morphologies despite changes occurring in the B cell germinal center. Network analysis, which we used to visualize the average distances between cell clusters and cell cluster abundance across the two TLS morphologies, demonstrated similar changes to the two B cell clusters (B_BCL6^high^ and B_BCL6^low^) occupying the germinal center and preservation of spatial relationships between mature dendritic cells, Tph, FOXP3^+^ T cells, proliferating T and B cells, and GZMB^+^ T cells is mature and involuted TLS (**Fig. 3h**). Overall, these neighborhood and network analyses suggested that while the B cell germinal center appeared to undergo dissolution in involuted TLS, the T cell zone was preserved.

Finally, evaluation of individual marker expression by cluster supported these observations regarding persistence of the T cell zone (**Fig. 3i**). In the mature dendritic cell cluster, expression of CCR7, HLADR, and CD86 were significantly increased, implying ongoing antigen presentation in these structures, and both the Tph and Tc clusters demonstrated increased expression of markers of antigen experience, including CD45RO, CD25, PD1, and TOX expression While the precise role of TOX in T peripheral helper cells is not established, TOX2 has previously been shown to be involved in the establishment of durable GC Tfh memory.^40^ Taken together, these data suggest that the involuted morphology may be a site of persistent antigen presentation by mature dendritic cells, which drive the formation of antigen-experienced memory T cell populations.

### Expanded T cell clonotypes are shared across TLS within a tumor, while B cell repertoires of individual TLS are highly distinct

Based on these data, we next sought to determine whether there were differences in the T and B cell repertoires of TLS of these two morphologies. We microdissected 38 individual TLS (32 mature and 6 involuted) from 7 treated tumors and performed bulk sequencing using the Adaptive ImmunoSEQ TCRβ and IGH assays (**Fig. 4a**, **Extended Data Fig. 7a, Extended Data Table 9**). After filtering to remove repertoires with low counts, 35 TCRβ repertoires and 32 IGH repertoires were analyzed. Across all samples, the repertoire size was variable with a mean total TCRβ clonotypes of 7171±8472 (**Extended Data Fig. 7b**). Singleton clonotypes comprised 68.7±13.4% of the TCRβ repertoire in all TLS sampled. In mature TLS, singleton clonotypes comprised 72.02±9.83% of the T cell repertoire, while in involuted TLS the singleton compartment constituted 48.98±16.1%. Across TLS microdissected from the same tumors, a mean of 32.3±12.3% of unique TCRβ clonotypes could be identified in other TLS from the same tumor. TCRβ clonotypes identified in all TLS from the same tumor were highly expanded, while those found in only one TLS were predominantly singletons (**Fig. 4b-c and Extended Data Fig. 7c-h**), suggesting a high degree of T cell trafficking as well as significant local T cell repertoire diversity at each individual TLS.

**Fig. 4.**
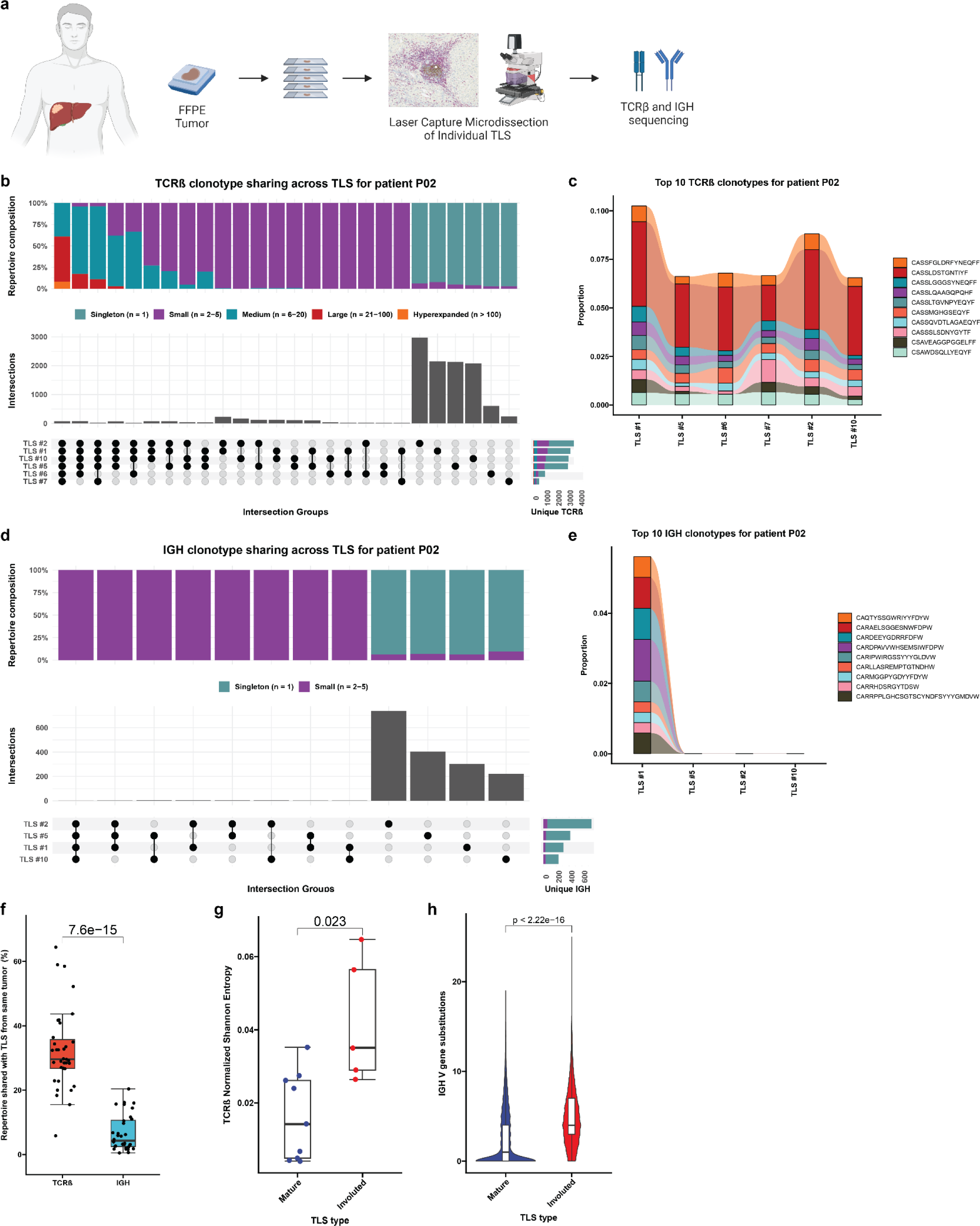
Expanded T cell clonotypes are shared across TLS within a tumor, while B cell repertoires of individual TLS are highly distinct. **a**, Workflow for T and B cell repertoire profiling of microdissected TLS (*n* = 30 mature and 5 involuted) from 7 patients. **b** and **d,** Upset plots showing overlap in unique TCRβ (**b**) and IGH (**d)** clonotypes across microdissected TLS from the same patient (P02). Barplots in gray and annotation row indicate distinct groups of clonotypes shared between different TLS. Top stacked barplots indicate composition of groups according to clonal expansion. Bottom right stacked barplots indicate total number of unique TCRβ or IGH clonotypes identified at each TLS according to degree of clonal expansion. **c** and **e,** Alluvial plots tracking the top 10 TCRβ (**c**) or IGH (**e**) clonotypes from TLS # 1 of patient P02 across all TLS microdissected from the patient’s tumor. **f,** Box-and-whisker plot comparing the percentage of the TCRβ or IGH repertoire of each TLS that is shared with other TLS from the same tumor. **g,** Box-and-whisker plots comparing TCRβ clonality (as determined by Normalized Shannon Entropy) in mature and involuted TLS microdissected from patients P12, OT1, and OT6. Each point represents the TCRβ of an individual TLS. **h,** Violin plots comparing number of somatic hypermutations in IGH of mature and involuted TLS microdissected from patients P12, OT1, and OT6. Individual data points (not shown) represent individual IGH sequences. Statistical significance was determined by two-tailed t test (**f-h**).

Across all microdissected TLS, the mean total number of IGH clonotypes was 922±1188 (**Extended Data Fig. 8a**). Singleton clonotypes comprised 95.3±3.9% of IGH repertoire of all TLS sampled. In mature TLS, singleton clonotypes comprised 95.7%±4.0 of the IGH repertoire, while in involuted TLS the singleton compartment constituted 93.4±3.1%. IGH repertoire sharing was significant lower across TLS microdissected from the same tumor (P = 7.6e-15), with only 6.7±5.6% of unique IGH clonotypes of each TLS detected in other TLS from the same tumor (**Fig. 4d-f** and **Extended Figure 7b-g**). These B cell repertoire characteristics are consistent with highly distinct, independent germinal center reactions.

In three patients (P12, OT1, and OT6) in which mature and involuted TLS were present in the same tissue block, we compared the immune repertoires of these two morphologies. TCRβ clonality was significantly increased in mature TLS compared to involuted TLS (P = 0.023) (**Fig. 4g**), although this difference was primarily observed in a single patient OT6 (**Extended Data Fig. 7i**). No difference was observed in IGH clonality (**Extended Data Fig. 7g**), but the IGH repertoire of involuted TLS did demonstrate a significantly higher number of V gene substitutions, a surrogate for somatic hypermutation (P < 2.22e-16) (**Fig. 4h** and **Extended Fig. 8i**). Taken together, these comparisons suggest that B cell populations in involuted TLS have undergone greater antigen-driven positive selection, consistent with a late-stage germinal center, and that there is associated T cell repertoire contraction and clonal expansion at these structures.

Given the extensive sharing of expanded T cell clonotypes observed within TLS from the same tumor, we also evaluated the peripheral blood to determine the extent of T cell trafficking between TLS and peripheral blood. We performed TCRβ sequencing of pre- and post-treatment peripheral blood mononuclear cells from 5 of the 7 patients whose TLS were microdissected. In TLS from these 5 patients, a mean of 44.0±8.4% of unique TCRβ clonotypes and 52.7±8.5% of total clonotypes in TLS were also identified in post-treatment peripheral blood. Similar overlap was observed between TLS repertoires and pre-treatment peripheral TCRβ repertoires, where a mean of 40.4±11.5% of unique TCRβ clonotypes and 48.7±11.0% total clonotypes in TLS were identified in pre-treatment peripheral blood (**Extended Data Fig. 9a-b**). In 3 of the 5 patients, 13 unique TCRβ were significantly expanded after neoadjuvant treatment and 9/13 (69.2%) were detected in at least one TLS (**Extended Data Table 10**). Together, these data provide evidence for a high degree of overlap between T cells within TLS and T cells in the peripheral blood.

### The top expanded T cell clonotypes in mature TLS are cytotoxic granzyme K and granzyme B-expressing CD8^+^ T cells

To further characterize T and B cell populations identified in TLS, we performed single cell RNA/TCR/BCR sequencing of post-treatment peripheral blood from the 7 patients from whose tumors TLS were microdissected. Sequencing of tumor infiltrating lymphocytes (TIL) was also performed for all 7 patients, but in only one sample (patient OT6) was sequencing data of sufficient quality for further analysis. Peripheral blood and TIL samples were processed by fluorescence-activated cell sorting (FACS) after labeling with antibodies to CD3 and CD19. After pre-processing and filtering to remove low quality sequencing data, 28,694 single cells were identified in the peripheral blood and 620 in the TIL. After performing preliminary cluster annotation using a reference annotated dataset, we attempted to match the TCRβ and IGH CDR3 amino acid sequences identified in microdissected TLS with sequences identified in the single cell dataset. TCRβ in the microdissection and single cell datasets were successfully matched (described below), but no matching IGH were identified between the bulk sequencing performed on microdissected TLS and single cell sequencing data, therefore B cells were excluded from subsequent analysis.

The resultant 23,172 T cells in the post-treatment peripheral blood samples and 562 T cells in the TIL of patient OT6 were clustered into 16 distinct cell clusters based on expression of canonical genes associated with specific T cell subsets. The CD4 compartment of the single cell dataset was divided into a naïve CD4^+^ T cell cluster (CD4 Naïve) expressing high levels of *CCR7* and *LEF1*; a CD4 Naïve-like cluster (CD4 Naïve-like) characterized by expression of *CCR7 and TCF7*; a CD4 T central memory cluster (CD4 TCM) with high expression of *LTB and S100A4*; a CD4 T peripheral helper cluster (Tph) characterized by low expression of *CXCR5,* high expression of *CXCR3,* and high expression of ICOS; and two CD4 T effector memory clusters notable for high expression of granzyme K (CD4 TEM_GZMK) and granzyme B (CD4 TEM_GZMB). CD8 T cells were divided into the following clusters: a naïve cluster (CD8 Naïve) highly expressing *CD8B*, *CCR7*, *LEF1*; a CD8 T central memory cluster (CD8 TCM) with elevated expression of *CD8B and LINC02446*; two CD8 T effector memory clusters distinguished by high expression of granzyme K (CD8 TEM_GZMK) and high expression of granzyme B (CD8 TEM_GZMB); and a CD8 tissue resident memory-like cluster (CD8 TRM) with increased expression of *NR4A2*, *DUSP2*, and *ZNF683.* In addition, we identified a CD4 regulatory T cell cluster (Treg) with high expression of *FOXP3* and *RTKN2*; an NK-T cell cluster (NK-T) highly expressing *PRF1* and *GZMB*; a double negative T cell cluster (dnT) with high expression of *SYNE* and *MALAT1*; a gamma delta T cells (gdT) with high expression of *TRDV2* and *TRGV9*; and mucosal invariant T cells cluster (MAIT) highly expressing *KLRB1* and *SLC4A10* (**Fig. 5a-c** and **Extended Data Fig. 10a-b**). In the TIL of patient OT6, 11 of 16 clusters were identified: CD4 Naïve-like, CD4 TCM, CD4 Tph, CD4 TEM_GZMK, CD8 TCM, CD8 TEM_GZMK, CD8 TEM_GZMB, CD8 TRM, Treg, dnT, and MAIT **(Extended Data Fig. 11a-c).**

**Fig. 5.**
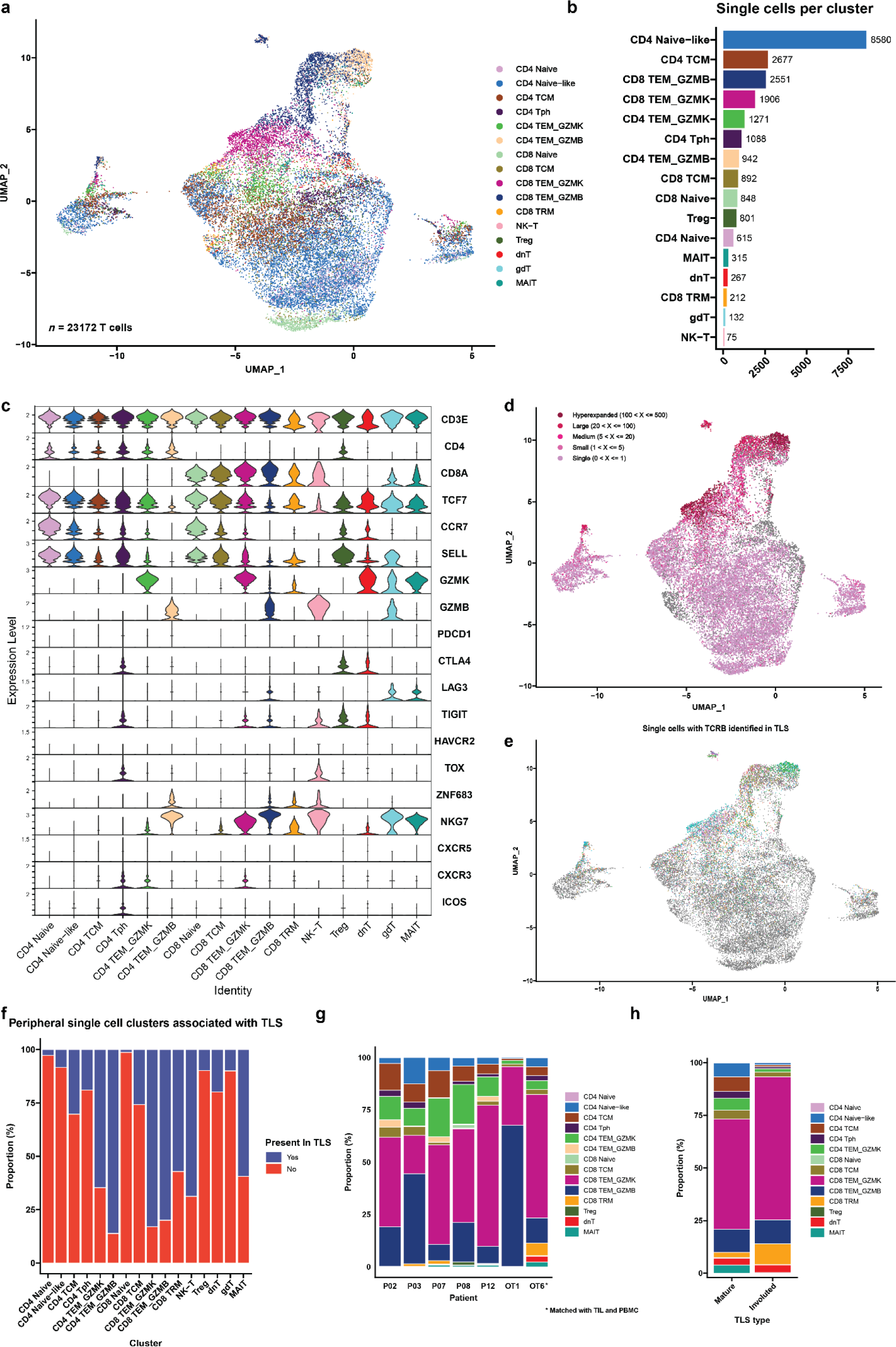
Cytotoxic granzyme K and granzyme B-expressing CD8 T cells are highly represented in TLS. **a**, Uniform Manifold Approximation and Projection (UMAP) of 23,172 T cells identified by single cell RNA/TCR/BCR sequencing of CD3^+^CD19^+^ FACS-sorted peripheral blood from HCC patients treated with neoadjuvant ICB (*n* = 7). **b,** Barplot showing number of single cells per cluster. **c,** Violin plots showing expression of subset specific marker genes across clusters. **d-e,** UMAPs showing clonality of single cells with an associated T cell receptor sequence (**d**) and single cells with a TCRβ identified in microdissected TLS (**e**). **f,** Stacked barplot showing proportion of each single cell cluster identified in TLS. **g,** Inferred transcriptional phenotype of TCRβ clonotypes in microdissected TLS with a matching TCRβ in single cell sequencing of post-treatment peripheral blood (*n* = 7) or tumor infiltrating lymphocytes (*n* = 1). **h,** Inferred transcriptional phenotype of TCRβ clonotypes in mature and resolving TLS of patient OT6.

19,546/23,172 (84.3%) single cells in the peripheral blood dataset and 346/562 (61.6%) cells in the TIL had a partial or completely sequenced TCRαβ chain identified by single cell TCR sequencing, of which there were 15,016 and 256 unique TCRs, respectively (**Extended Data Table 11**). In the peripheral blood single cell dataset, clonal expansion was most strongly associated with the CD4 TEM_GZMB (Odds Ratio 31.48, P < 0.001) and CD8 TEM_GZMB clusters (Odds Ratio 17.99, P < 0.001) by Fisher’s Exact test (**Fig. 5d** and **Extended Data Table 12**). In TIL, clonal expansion was most strongly associated with the CD8 TEM_GZMB (Odds Ratio 3.49, P = 0.003), CD8 TRM (Odds Ratio 2.57, P = 0.041), and CD8 TEM_GZMK (Odds Ratio 1.94, P = 0.013) clusters (**Extended Data Fig. 11d and Extended Data Table 12).**

T cells belonging to the CD8 TEM_GZMK cluster were notable for increased expression of the gene associated with cytotoxicity, including Granzyme K (*GZMK*) and the chemokine ligand CCL5 (*CCL5*) and decreased expression of Granulysin (*GNLY*). The CD8 TEM_GZMB clusters also demonstrated hallmarks of cytotoxicity, including elevated expression of granzyme B (*GZMB*) and granzyme H (*GZMH*), as well as elevated expression of perforin (*PRF1*), and *GNLY* (**Extended Data Fig. 10c-d**).^41^ *GZMK* expression was low in the latter cluster. These two transcriptional phenotypes were consistent with T progenitor exhausted and cytotoxic/terminally differentiated CD8 T cell states, respectively, which have been previously identified in the peripheral blood and tumors of patients treated with ICB.^41^ Consistent with this identity, in the TIL from patient OT6, both *GZMK* and *GZMB* expressing CD8 clusters showed increased expression of multiple T cell exhaustion markers, including *PDCD1*, *CTLA4*, *LAG3*, *TIGIT*, and *TOX*, which were more highly expressed in CD8 TEM_GZMB compared to CD8 TEM_GZMK. *NKG7* and *CCL5*, which are associated with cytotoxic CD8 T cells, were also increased in both clusters, with higher expression in the GZMB high cluster (**Extended Data Fig 11h-i**). Notably, both clusters in the single cell TIL from patient OT6 demonstrated elevated expression of the B cell chemoattractant *CXCL13*, which has been associated with tumor-specific T cells in single cell sequencing studies of TIL from patients treated with ICB and is involved in the formation of TLS.^42,43^

No CD4^+^ T cell cluster was observed in the single cell data that displayed a transcriptional phenotype consistent with a CD4^+^ T follicular helper (Tfh) population, which are defined by high expression of CXCR5, CXCR3, and ICOS. Instead, we identified a CD4^+^ T cell cluster consistent with the T peripheral helper cluster identified by imaging mass cytometry. In the peripheral blood, cells belonging to this cluster demonstrated low CXCR5 expression, increased CXCR3 expression, and elevated expression of CTLA4, TIGIT, and TOX (**Extended Data Fig. 10e**). Cells belonging to this cluster in the single cell TIL demonstrated increased expression of *CXCL13*, *ICOS*, *PD1*, *MAF*, *TOX* and high expression of multiple exhaustion markers including *CTLA4*, *LAG3*, *TIGIT*, *HAVCR2*, and *TNFRSF18 (GITR)* (**Extended Data Fig 11j**). Based on these data and the imaging mass cytometry above, we conclude that Tph constitute a major CD4^+^ T cell subset found in TLS in the context of neoadjuvant immunotherapy.

Approximately one-third, or 6349/19546 (32.5%), of single cells with a TCRβ in the peripheral T cell dataset and 199/346 (77.7%) single cells with a TCRβ in the TIL were identified in at least one TLS (**Extended Data Table 11**). TCRβ identified in TLS were detected in all clusters of the peripheral and TIL single cell dataset. Among peripheral blood T cells, the clusters most strongly associated with TLS were CD4 TEM_GZMB (Odds Ratio 10.72, P < 0.001), CD8 TEM_GZMB (Odds Ratio 9.73, P < 0.001), and CD8 TEM_GZMK (Odds Ratio 9.46, P < 0.001) (**Fig. 5e-f** and **Extended Data Table 13**), suggesting that TLS specifically promote the trafficking of effector memory CD4^+^ and CD8^+^ T cell populations from the peripheral blood to tumor. In the TIL, no cluster was significantly correlated with TLS but the Treg cluster was inversely correlated with presence in TLS (Odds Ratio 0.22, P = 0.003) (**Extended Data Fig. 11e-f** and **Extended Data Table 13**). The proportion of all TCRβ identified in TLS that were matched to single cell data was low overall (2,908/135,909 unique clonotypes or 2.1%), but a higher proportion of clonotypes were successfully matched for the most expanded clones, with 369/1359 (27.2%) of the top 1% of TCRβ that had been identified in TLS being successfully matched to the single cell data, and 63/137 (46%) of the top 0.1% of TCRβ (**Extended Data Table 14**). Thus, this approach, while providing a limited view of singleton TCRβ identified in TLS, could be used to provide additional transcriptional information about expanded T cell populations trafficking to TLS.

We next used these data to infer the transcriptional phenotype of T cells trafficking through TLS. Since our single cell dataset contained peripheral blood and TIL data, we first evaluated the correlation between peripheral blood and TIL phenotypes, and the reliability of using data from single cell sequencing from one compartment to infer the properties of single cells with the same TCRβ in the other compartment. To carry this out, we examined the correlation between single cell cluster identity for TCRβ which were found in both peripheral blood and TIL in patient OT6. In total, 16 unique TCRβ sequences were present in both peripheral blood and TIL (**Extended Data Fig. 11a**). 7/16 TCRβ clonotypes had the same single cell identity for all cells with the same TCRβ in the peripheral blood and TIL, and an additional 5/16 TCRβ had the same cluster identity assigned to at least half of the cells in both peripheral blood and TIL. In only 4/16 TCRβ were the cluster identities of cells with the same TCRβ entirely discordant (**Extended Data Fig. 11b).** Based on these data, we concluded that the peripheral blood transcriptional phenotype closely recapitulates the cluster assignment of TIL, and thus both identities may be used to determine a putative phenotype for T cells identified in TLS by TCRβ sequencing. These findings were consistent with previous work demonstrating that in circulating TILs, gene signatures of effector functions, but not terminal exhaustion, reflect those observed in the tumor.^44^

Across all 7 patients, the majority of T cells identified by matching of the TCRβ were GZMK and GZMB expressing CD8 T effector memory cells, but we also observed CD4 TEM_GZMK, CD4 CTL, and CD4 T peripheral helper clusters among the putative phenotypes of T cells trafficking through TLS (**Fig. 5g**). Notably, in the involuted TLS from the tumor of patient OT6, where we had previously noted significant increase in clonality relative to mature TLS, clonal expansion was greatest in the of CD8 TEM_GZMK, CD8 TEM_GZMB, and CD8 TRM clusters (**Fig. 5h** and **Extended Data Fig. 11k**). Overall, these data provide single cell resolution to the top expanded clonotypes in TLS and show that highly expanded T cell populations in TLS are CD8^+^ T cell effector memory, which may undergo clonal expansion and repertoire contraction in concert with expansion of resident memory populations in areas of tumor regression.

## DISCUSSION

Neoadjuvant immunotherapy aims to use the primary tumor as a source of antigens to enhance antitumor immunity and prevent cancer recurrence after surgery.^4^ Preclinical and clinical data suggest that this approach induces more durable immunologic memory than adjuvant immunotherapy alone,^3^ but the mechanism by which this occurs and the contribution of TLS to this process are not well understood. The data presented here show that in HCC treated with neoadjuvant immunotherapy, intratumoral TLS are associated with superior pathologic responses and improved relapse free survival. These findings are consistent with data reported in other solid tumors treated with ICB,^9–12^ as well as studies of the prognostic significance of intratumoral TLS in treatment-naïve early stage HCC treated with surgical resection.^16^ In tumors with high TLS density and significant regression of the tumor, we further identified an involuted morphology of TLS in areas of nonviable tumor whose location, histologic, and immunologic features, and similarity to late stage germinal centers observed in murine secondary lymphoid organs,^26^ are consistent with a terminal stage of the TLS life cycle. Using laser capture microdissection, bulk immune repertoire sequencing, and matched single cell sequencing, we identify and characterize expanded T cell populations trafficking through TLS and find evidence for significant immune repertoire changes associated with TLS dissolution.

While TLS are thought to mature from a loosely organized lymphoid aggregate to a CD21^+^ primary follicle and reach full maturity as a CD21^+^CD23^+^ secondary follicle,^16,45,46^ which have distinct T and B cell zones,^25^ the circumstances of TLS resolution are not known.^47^ These data suggest that TLS dissolution may be driven by elimination of tumor and may occur dyssynchronously, with dissolution of the B cell germinal center accompanied by persistence of a T cell zone enriched for interactions between DCLAMP^+^CCR7^+^HLADR^+^ mature dendritic cells and CD4^+^ and CD8^+^ T cells. Furthermore, the changes observed in T cell repertoire at these structures, including increase in clonality and expansion of cytotoxic and tissue resident memory-like CD8^+^ T cell clonotypes, suggest that late-stage TLS may play a functional role in supporting the contraction and memory phase of the intratumoral adaptive immune response through persistent antigen presentation in the T cell zone (**Fig. 6**). Such a role would also be consistent with recent data suggesting that tonic antigenic stimulation drives programs of T cell residency in tumors, and would identify a specific place where such interactions may occur.^48^

**Fig. 6.**
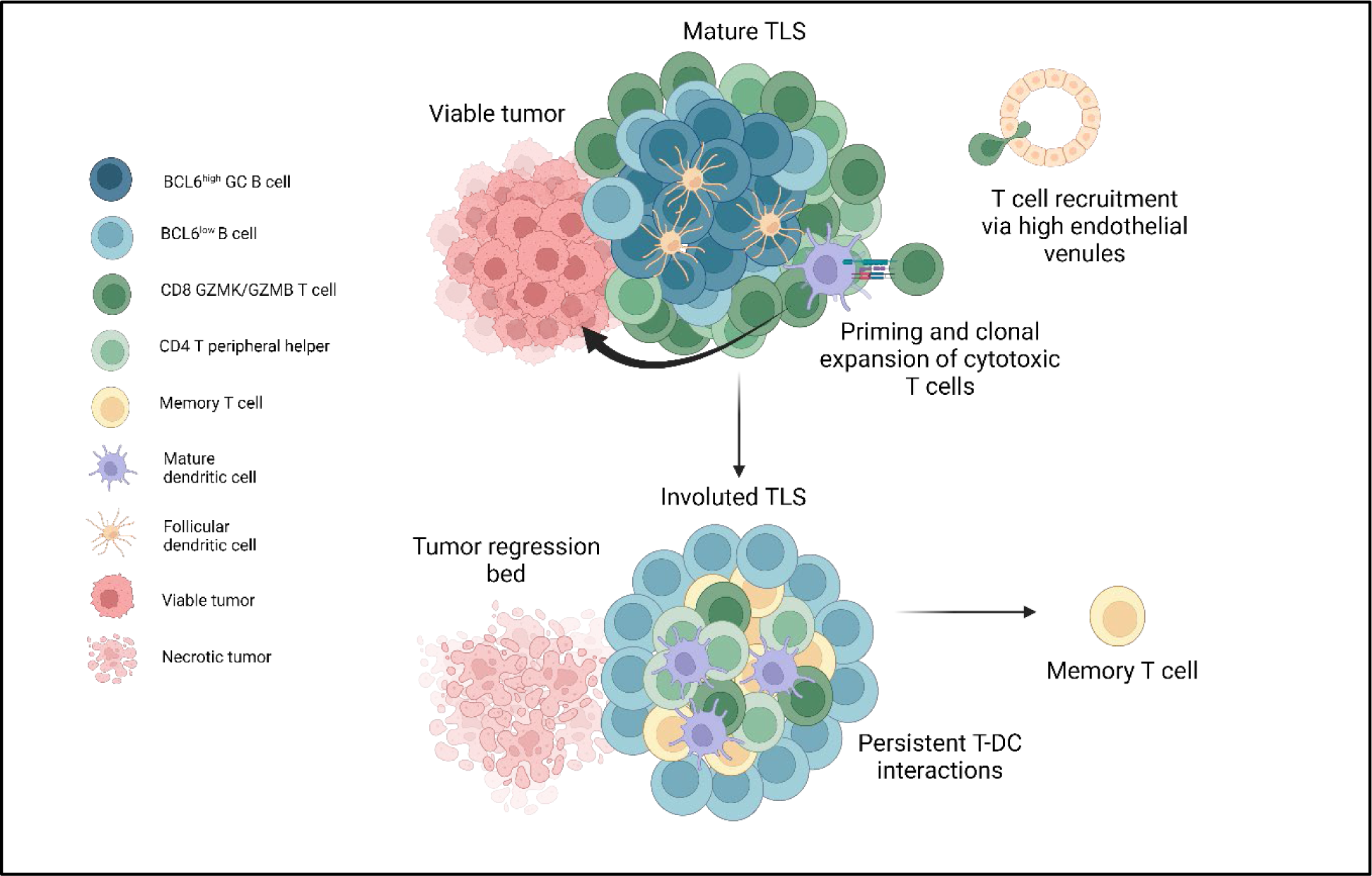
TLS structure and function in viable tumor and tumor regression bed in tumors treated with neoadjuvant checkpoint blockade. Mature TLS in viable tumor display a highly organized germinal center with close interactions between germinal center B cells and CD21^+^ follicular dendritic cells, a T cell zone characterized by CD4^+^ T peripheral helper cells in close proximity to mature dendritic cells, and cytotoxic CD8^+^ T cells trafficking to the tumor via high endothelial venules. In areas of tumor regression, an involuted TLS morphology is found which displays dissolution of the B cell germinal center and persistence of Tph-DC interactions in the T cell zone, increased T cell memory marker expression, and clonal expansion of cytotoxic and tissue resident memory CD8^+^ T cells.

Notably, in neoadjuvant treated tumors we did not detect a CXCR5^+^CXCR3^+^CD4^+^ T follicular helper population, which have been reported in tumor associated TLS.^49,50^ Rather, in both the imaging mass cytometry analysis and single cell datasets we identified a CXCR5^-^CXCR3^+^ CD4^+^ T cell population, which expressed CXCL13 in the single cell TIL of patient OT6 and was also detectable in post-treatment peripheral blood single cell sequencing. CXCR5^-^CXCL13-producing CD4^+^ T cells have been identified in untreated human breast cancer under the name TFHX13^51,52^ and in autoimmune disease, where they are termed CD4^+^ T peripheral helper cells and have been shown to provide help to B cells in an IL-21 dependent manner.^34,35,53^ This population was present in both the mature and involuted morphologies observed in these patients. Additional studies are required to determine whether this population of CD4^+^ T cells provides help to B cells in this treatment setting, and to determine their role in involuted TLS, where they were found in association with mature dendritic cells.

We recognize several limitations of the findings reported here. First, samples were obtained from a single institution and may not represent the full diversity of HCC etiologies and subtypes. Additionally, while the untreated cohort in this study was similar to the treatment cohort in age, sex, and HCC etiology and received treatment at the same primary institution, we cannot exclude the possibility that the different pathologic findings between the untreated and neoadjuvant cohorts arose as a consequence of differences in the patient populations rather than treatment status. We infer the transcriptional phenotype of T cells infiltrating TLS based on matching TCRs identified by microdissection with peripheral T cells and/or TIL subjected to single cell sequencing. In our data, we demonstrate a correlation between the cluster identity for 16 TCRs shared between the peripheral blood and TIL of patient OT6, and others have reported correlation between gene signatures of effector functions in circulating TILs and tumor;^44^ however, it is possible that the phenotype of these T cells are not fully conserved outside of TLS. Finally, our analyses throughout are limited by small sample size. However, clinical samples from neoadjuvant studies are rare, and as we have shown here even small samples can provide important insights into the constituent immune populations in tumors arising in human subjects.

Finally, these data raise several important questions which future studies should address. First, the role of FOXP3^+^ T follicular regulatory cells, which have been shown to regulate germinal center reactions,^54,55^ is not completely understood in this context and this population could not be resolved in our imaging mass cytometry analysis. Second, while dispersion of the B cell follicle was associated with attenuation of the PDPN^+^ fibroblastic reticular cell and the CD21^+^CD23^+^ follicular dendritic cell network in our data, other changes occurring in the fibroblast populations involved in this process remain unclear and it is not known what permits retention of mature dendritic cells within the T cell zone. Third, since all tumors in which involuted TLS were observed were treated with neoadjuvant ICB and these structures were not seen in untreated tumors, the contribution of therapy to their formation cannot be established from our data. In murine studies, both the PD-1 and CTLA-4 pathways have been shown to be involved in regulation of T follicular regulatory populations,^56–58^ suggesting that therapeutic blockade may affect the dynamics of germinal center formation and dissolution. Nonetheless, these data shed light on the circumstances of TLS resolution and suggest that this terminal stage, about which nothing was previously known, may play a functional role in the formation of intratumoral T cell memory after elimination of viable tumor.

## METHODS

### Study design

The aim of this study was to characterize tertiary lymphoid structures (TLS) in patients with hepatocellular carcinoma (HCC) treated with neoadjuvant ICB-based therapy prior to surgical resection of the primary tumors. To understand the clinical significance of TLS, we analyzed TLS density in treated patients and untreated controls and correlated TLS density in treated patients against pathologic response and post-surgical clinical outcomes. We performed bulk RNA sequencing of tumors with high and low TLS density after neoadjuvant treatment to understand the gene expression programs of tumors with high TLS density. We then characterized the morphological and functional properties using bulk immune repertoire sequencing of TCRβ and IGH, imaging mass cytometry, matched single cell TCR and RNA sequencing of peripheral blood and tumor infiltrating lymphocytes.

### Patient identification and data collection

Patients were identified for inclusion in this study if they received surgical resection for locally advanced, non-metastatic hepatocellular carcinoma after neoadjuvant ICB-based therapy between October 1, 2019 and January 31, 2022 at the Johns Hopkins Sidney Kimmel Cancer Center. Retrospective chart review was performed to collect clinical data from the electronic medical record regarding age at surgery, sex, date of resection, HCC etiology, histologic grade of tumor, neoadjuvant treatment, relapse free survival, and overall survival. A cohort of untreated control patients who had undergone surgical resection for HCC without prior systemic treatment were also identified via a search of the electronic medical record. Review of the electronic medical record was performed to confirm absence of prior systemic treatment. For both cohorts, histologic grade was based on pathologic assessment at the time of resection if there was discordance with grade reported for pre-treatment biopsy. Patients in both cohorts were excluded from analysis if there was evidence of active hepatitis B (defined by a positive HBsAg or detectable HBV DNA) prior to surgery. Patients were excluded from the control group if the etiology of their HCC was not represented in the treatment group (e.g. hepatic adenoma and hereditary hemochromatosis). This study was conducted in accordance with the Declaration of Helsinki and was approved by the Johns Hopkins University Institutional Review Board (IRB00149350, IRB00138853, NA_00085595). Informed consent or waiver of consent was obtained from all patients. Treated patients identified with the letter P were accrued as participants in the phase I clinical trial NCT03299946.^8^

### Histopathologic assessment of TLS density and pathologic response

Evaluation of pathologic response was performed by a hepatopathologist (RA). Pathologic response designations were assigned according to percent residual viable tumor in surgically resected tumors. Complete response (CR) was defined as 0% residual viable tumor, major pathologic response (MPR) as less than 10% residual viable tumor, partial pathologic response (pPR) as 10-90% residual viable tumor, and non-response (NR) as >90% residual viable tumor.^17^ 12 of the 19 patients had previously undergone assignment of pathologic response according binary categorization of major or complete pathologic response versus non-response,^8^ and for this group non-responders were categorized as NR or pPR as described above. To determine TLS density, formalin fixed paraffin embedded (FFPE) tumors were sectioned, mounted on glass slides, and stained with anti-CD20 antibody as described below. Whole slide images were obtained at 0.49 µm per pixel using the Hamamatsu NanoZoomer. The presence of CD20 positivity was determined by digital image analysis software (HALO v3.0.311 Indica Labs), with TLS defined as CD20 positive cell aggregates greater than 150μm in diameter located among tumor cells or at the invasive margin in areas of viable and nonviable tumor. TLS density was determined by calculating the number of TLS per mm^2^ of viable and nonviable tumor. TLS were classified as peritumoral if they were found within 200 μm of the interface between normal adjacent parenchyma and tumor and intratumoral if they were found within the tumor or tumor regression bed.

### Survival analyses

The Kaplan–Meier method was used to estimate relapse-free survival (PFS) and overall survival (OS). Relapse free survival (RFS) was defined as the time from surgical resection to radiographic relapse. Overall survival (OS) was defined as the time from surgical resection to death from any cause. If a patient was not known to have had either event, RFS and OS were censored at the last date of known healthcare contact. RFS and OS analyses were limited to patients treated with neoadjuvant therapy and were not performed in the untreated controls due to limited follow up in this cohort. Survival analyses using the Kaplan-Meier method were performed using the R package survminer. Bayesian information criterion (BIC) analysis was performed using the R package stats. A linear regression model was used to evaluate the effect of each marker, dichotomized by the mean, as a predictor of each distance measure. For each binary outcome, logistic regression was employed, with each marker treated as continuous. A meaningful difference in BIC between the two models is 2 at a minimum, and a difference between 5-10 and above 10 is considered to be strong and very strong, respectively.^59^

### Immunohistochemistry

Automated single and dual staining was performed on the Leica Bond RX (Leica Biosystems). Single staining for CD20 was employed for determination of TLS density. Dual staining for CD3 and CD21, CD8 and CD4, Ki67 and CD20 was performed prior to laser capture microdissection of TLS. Slides were baked and dewaxed online followed by antigen retrieval for 20 min at 100°C. Endogenous peroxidase was blocked using Peroxidase block (Refine Kit) followed by Protein block (X090930-2, Agilent Technologies Inc., Santa Clara, CA). Primary antibodies were applied at room temperature. Detection was performed using the Bond Polymer Refine Kit (DS9800, Leica Biosystems). For dual staining, a second round of antigen retrieval was performed for 20 min at 95°C followed by application of a second primary antibody. Detection of the second primary antibody was performed using the Bond Polymer Red Refine Kit (DS9390, Leica Biosystems). Slides were counterstained, baked and coverslipped using Ecomount (5082832, Biocare Medical, Walnut Creek, CA). Antigen retrieval buffers and concentrations of all antibodies are listed in **Extended Data Table 15**. Antibodies were diluted to appropriate working concentration using Antibody Diluent (S302283-2, Agilent Technologies Inc).

### Bulk RNA sequencing and TCRβ/BCR immune repertoire profiling of FFPE tumor

RNA was extracted from FFPE tumor from the treatment cohort and sequenced using the commercial platform ImmunoID NeXT with 200 million paired end reads (150 base pair). Reads were aligned in accordance with the Personalis Cancer RNA pipeline and transcript per million (TPM) values were extracted.^60^ Bulk RNA sequencing was performed on 14 tumors in two batches. No batch correction was applied due to lack of clear batch-to-batch differences by principal component analysis. 2 samples were excluded due to poor sequencing depth, as defined by a median of the log2 transformed count data being equal to 0 for those samples. The remaining 12 samples were filtered to include only genes for which the sum of raw counts across all samples was greater than 1. Variance stabilizing transformation was performed on the resultant data and differentially expressed genes were identified using DESeq2.^61^ Genes with an adjusted P value of < 0.05, and a minimum log2 fold change of 1 were considered differentially expressed. Pathway analysis was performed using the R package fsgsea to identify biologically enriched pathways from the MSigDB hallmark gene sets.^62,63^ For pathway analyses, adjusted P values of < 0.05 were considered statistically significant. TCRβ and BCR repertoire profiling was performed using the ImmunoID NeXT transcriptome, which provides augmented (approximately a 100x increase over a standard transcriptome) coverage of TCRβ and BCR.^60,64^ Clones were identified using MiXCR^65^ and repertoire analysis was performed using the R package immunarch.^66^ Clonality was calculated as 1-Shannon’s equitability^67^ with clonality values ranging from 0-1, with 0 indicating equal representation of all clones within a repertoire and 1 being a repertoire consisting of only one clone.

### IMC staining and acquisition

IMC Staining was done as previously described.^8,68^ Briefly, formalin-fixed paraffin-embedded (FFPE) resected liver tissue sections were baked, deparaffinized in xylene, then rehydrated in an alcohol gradient. Slides were incubated in Antigen Retrieval Agent pH 9 (Agilent PN S2367) at 96 °C for 1 hour then blocked with 3% BSA in PBS for 45 min at room temperature followed by overnight staining at 4°C with the antibody cocktail. Antibodies, metal isotopes, and their titrations are listed in **Extended Data Table 8**. Images were acquired using a Hyperion Imaging System (Standard BioTools) at the Johns Hopkins Mass Cytometry Facility. Upon image acquisition, representative images were visualized and generated through MCD™ Viewer (Standard BioTools).

### IMC data analysis

Images were segmented into a single-cell dataset using the publicly available software pipeline based on CellProfiler, ilastik, and HistoCAT.^69–72^ Since multiple images contained more than one TLS, images were subset for distinct TLS regions by manual gating using FlowJo^TM^v10.9.0 Software (BD Life Sciences), which identified the xy coordinates of cells belonging to distinct lymphoid aggregates (**Extended Data Fig. 6b-c**). This resulted in 38 unique TLS matching either the mature (*n* = 20) or involuted morphology (*n* = 18). The resulting 61,371 single cells were clustered using FlowSOM^73^ into metaclusters, which were manually annotated into final cell types. Density of each cell type was determined by calculating the number of cells per unit area as determined by ImageJ v1.53.^74^ For network visualization, the mean distance between each cell type was computed and visualized using the R package qgraph.^75^ Neighborhood analysis was performed by using data generated by HistoCAT summarizing the top neighboring cell types for every cell type.

### Laser capture microdissection and TCRβ/IGH sequencing of microdissected TLS

10-14 µm serial tissue sections were obtained from formalin-fixed paraffin embedded (FFPE) tumor tissue blocks and mounted on UV activated PEN membrane glass slides (Applied Biosystems Cat. No. LCM0522) with additional 4-µm tissue sections cut every 150 µm for staining with hematoxylin and eosin (H&E) and dual IHC for CD3/CD21, CD8/CD4, and Ki67/CD20 as described above. Stained sections were scanned at 20x objective equivalent (0.49 µm pixel^−1^) on a digital slidescanner (Hamamatsu Nanozoomer) in advance of microdissection and annotated using NDP.view2 viewing software in order to identify areas for microdissection. On the day of microdissection, unstained tissue sections mounted on PEN membrane slides were deparaffinized using xylene and graded alcohol washes and stained with H&E. Laser capture microdissection of individual TLS was performed on the LMD 7000 system (Leica) and genomic DNA was extracted using the Qiagen QIAamp DNA FFPE Tissue Kit following the manufacturer’s protocol (Qiagen). DNA concentrations were quantified with a Qubit 4 Fluorometer using the Qubit dsDNA high sensitivity assay (Invitrogen). Sequencing of the TCRβ and IGH CDR3 regions was performed using the immunoSEQ platform (Adaptive Biotechnologies).^76,77^ TCRβ and IGH repertoire data were downloaded from the Adaptive ImmunoSEQ analyzer web interface after filtering to remove non-productive reads. After exclusion of repertoires with fewer than 500 TCRβ clones and 50 IGH clones, subsequent analysis was performed using the R package immunarch^66^ and the Python package Change-O,^78^ which was used to assign clonal families to IGH data. J gene, and greater than 90% identical CDR3 sequence according to nucleotide hamming distance. Clonality was calculated as described above using 1-Shannon’s equitability. To compare clonality across multiple TLS from the same tumor, we used the median clonality of 1000 iterations of downsampling to the number of productive CDR3 sequences in the smallest TCRβ or IGH repertoire for that patient.^79^

### Peripheral blood and fresh tumor collection and processing

Processing of peripheral and cryopreservation was completed as previously described.^8^ Fresh tumor tissue was diced with a sterile scalpel and dissociated in 0.1% collagenase in RPMI 1640 for 60 minutes at 37⁰C using the gentleMACS OctoDissociator (Miltenyi Biotec) according to the manufacturer’s instructions. Supernatant was collected and centrifuged at 1500 rpm for 10 minutes. Supernatant was removed and discarded, and the cell pellet was resuspended in ACK Lysing buffer (Quality Biological, cat# 118-156-721) and incubated at room temperature for 5 minutes before centrifugation at 1500 rpm for 10 minutes. Cells were resuspended in PBS, counted using a manual hematocytometer, and cryopreserved in 10% DMSO/AIM-V freezing media.

### Single cell RNA/TCR/BCR-sequencing

For all 7 patients from whose tumors TLS were microdissected, single cell sequencing was obtained for peripheral blood T and B cells isolated by Fluorescent Activated Cell Sorting (FACS). For 6 of 7 patients, the peripheral blood sample was obtained following completion of neoadjuvant ICB and prior to surgical resection; for 1 of the patients, the peripheral blood sample was drawn 4 weeks after resection. In addition, in the latter patient, single cell sequencing was performed for tumor infiltrating T and B cells isolated by FACS from tumor specimen. Cryopreserved PBMC and tumor suspension were thawed and washed with pre-warmed RPMI with 10% FBS. Cells were resuspended 0.04% BSA in PBS and stained with a viability marker (Zombie NIR, BioLegend) and Fc block (Biolegend, Cat. no. 422302) for 10 minutes at room temperature in the dark. Cells were then stained with antibodies against CD3 (FITC, clone HIT3a), for 20 minutes on ice and CD19 (PE/dazzle, clone SJ25C1) (**Extended Data Table 16**). After staining, viable CD3^+^ and CD19^+^ cells were sorted into 0.04% BSA in PBS using a BD FACS Aria II Cell Sorter at a 4:1 ratio. Sorted cells were counted and resuspended at a concentration of 1000 cells per μl. The single-cell library preparations for gene expression and V(D)J were performed with the Chromium Next GEM Single Cell 5’ GEM Kit v2 (10x Genomics) and Chromium Single Cell V(D)J Amplification Kit (human TCR) (10x Genomics), respectively. The cells were partitioned into nanoliter-scale Gel Beads in-emulsion (GEMs) and cells were barcoded. The cDNA synthesis and amplification was performed prior to sample split for the gene expression and for V(D)J libraries. Single cell libraries were sequenced on an Illumina NovaSeq instrument using 2 × 150- bp paired end sequencing. 5′ VDJ libraries were sequenced to a depth of 5,000 reads per cell. 5′ DGE libraries were sequenced to a depth of 50,000 reads per cell.

### Single cell data pre-processing, quality control, clustering and integration

Cell Ranger v6.1.2 was used to demultiplex FASTQ reads, perform sequence alignment to the GRCh38 transcriptome, and extract unique molecular identifier (UMI) barcodes. Single cell gene expression matrices were analyzed using the R package Seurat v4.1.1 as a single Seurat object. Cells were filtered to include only cells with less than 25% mitochondrial RNA content and between 200 and 4000 genes detected. For single-cell VDJ sequencing, only cells with full-length sequences were retained. Raw count data were normalized using the Seurat function SCTransform to normalize raw count data to a Gamma-Poisson Generalized Linear Model, perform variance stabilization, identify highly variable features, and scale features.^80,81^ Cells were projected into their first 50 principal components using the RunPCA function in Seurat, and further reduced into a 2-dimensional visualization space using the RunUMAP function. Initial cell cluster identification was performed using the Seurat function FindClusters at a resolution of 0.7. Initial cell type assignment was performed by reference mapping to the human PBMC dataset associated with the R package Azimuth.^82^ Cluster identities were then manually assigned by identification of differentially expressed genes using the MAST hurdle model as implemented in the Seurat FindAllMarkers function with a log fold change threshold of 0.25 and minimum fractional expression threshold of 0.25.^83^ Integration of single cell TCR-seq and BCR-seq data into the scRNA-seq data was performed using the R package scRepertoire.^84^ For each patient, TCRβ sequences identified in single cell data were compared against TCRβ identified in microdissected TLS to identify T cells present in TLS. In cases where single cells with the same TCR occupied multiple clusters, a putative transcriptional phenotype was assigned to a T cell in the TLS repertoire according to the most common T cell subset to which the single cells with the same TCR belonged. No matches were identified between IGH sequences identified by Adaptive sequencing and IGH sequences in the single cell dataset, and thus we excluded B cells in the single cell dataset from further analysis.

## Data availability

Bulk RNA-seq, single cell RNA/TCR-seq data, and imaging mass cytometry data from this study are deposited in dbGap under *** and the Gene Expression Omnibus (GEO) under accession number GSE ***. Bulk TCRβ and IGH data from microdissected TLS are available on the Adaptive ImmunoSEQ web analyzer portal at ***. All other relevant data are available from the corresponding authors upon request.

## Code availability

All custom code used to generate the results in this study has been deposited in a GitHub repository at https://github.com/FertigLab/HCCTLS.

## Acknowledgements

This work was supported by the Johns Hopkins SPORE in Gastrointestinal Cancer and the National Institutes of Health (NIH) U01CA253403 and U01CA212007 grants to M.Y. and E.J.F. Additional research support was provided by the Breeden-Adams Foundation and Conquer Cancer to M.Y.; the Johns Hopkins School of Medicine J. Mario Molina Physician Scientist Fund, Linda Rubin Fellowship, and Bernice Garchik Fellowship to D.S; and the Maryland Cancer Moonshot Research Grant to the Johns Hopkins Medical Institutions to D.J.Z. We thank Theinmozhi Arulraj for reviewing the final manuscript.

## Author contributions

D.S., M.Y., and E.J.F. conceived and designed this study. D.S., L.K., M.Y., and L.D. performed data analysis and interpreted results. Q.Z. and R.A. performed the pathologic review. D.S., K.M., Q.Z., and R.A. performed the histologic analysis. All authors assisted with the data analysis, provided valuable discussion, and reviewed and edited the final manuscript draft. D.S. and M.Y. wrote the manuscript with input from all the authors.

## EXTENDED DATA

**Extended Data Fig. 1.**
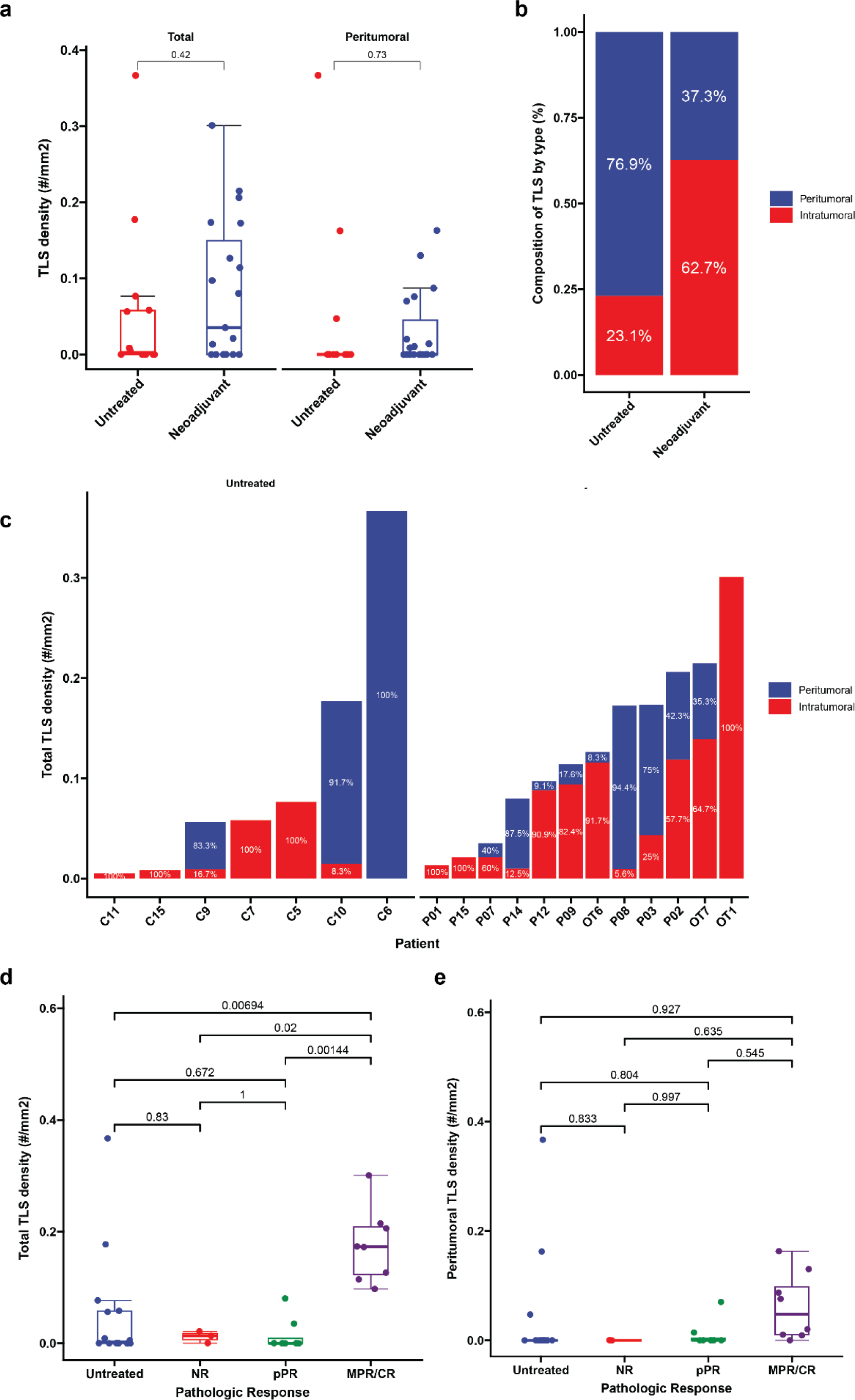
TLS density in HCC tumors treated with neoadjuvant ICB and untreated controls. **a**, Box-and-whisker plots showing total and peritumoral TLS density in patients with locally advanced HCC treated with neoadjuvant ICB (*n* = 19) and untreated controls (*n* = 14). **b-c,** Stacked barplots showing proportion of TLS comprised of peritumoral verus intratumoral TLS location neoadjuvant treated and untreated HCC tumors (**b**) and by patient (**c**). Labels indicate proportion of total TLS comprised of peritumoral or intratumoral TLS. In **c**, patients with no observed TLS are not shown. **d-e,** Box-and-whisker plots showing total (**d**) and peritumoral (**e**) TLS density in untreated (*n* = 14) and neoadjuvant treated tumors, divided according to pathologic response (*n* = 19). Statistical significance was determined by two-tailed t-test (**a**) and one-way ANOVA followed by Tukey’s honest significant difference (HSD) test (**d** and **e**).

**Extended Data Fig. 2.**
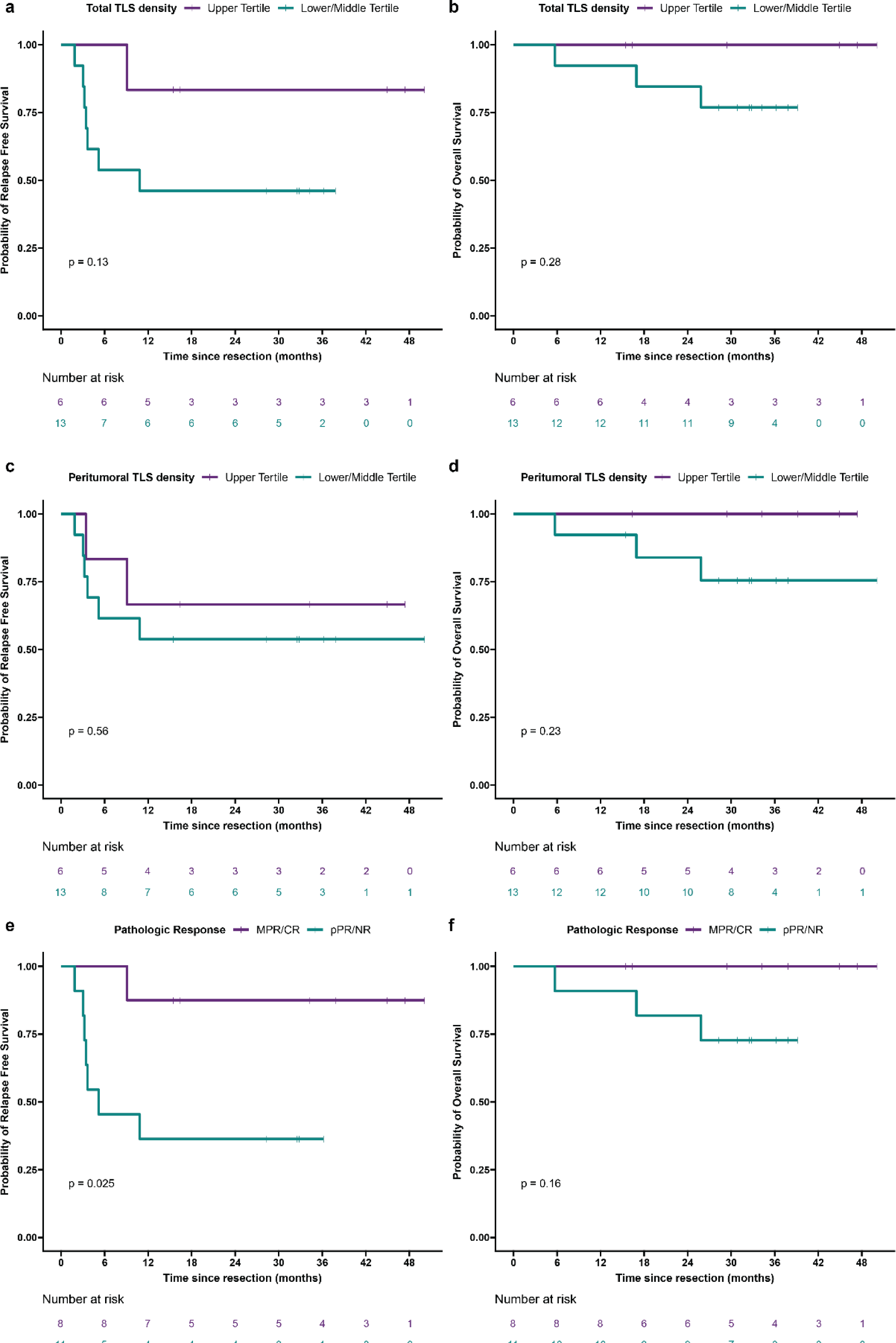
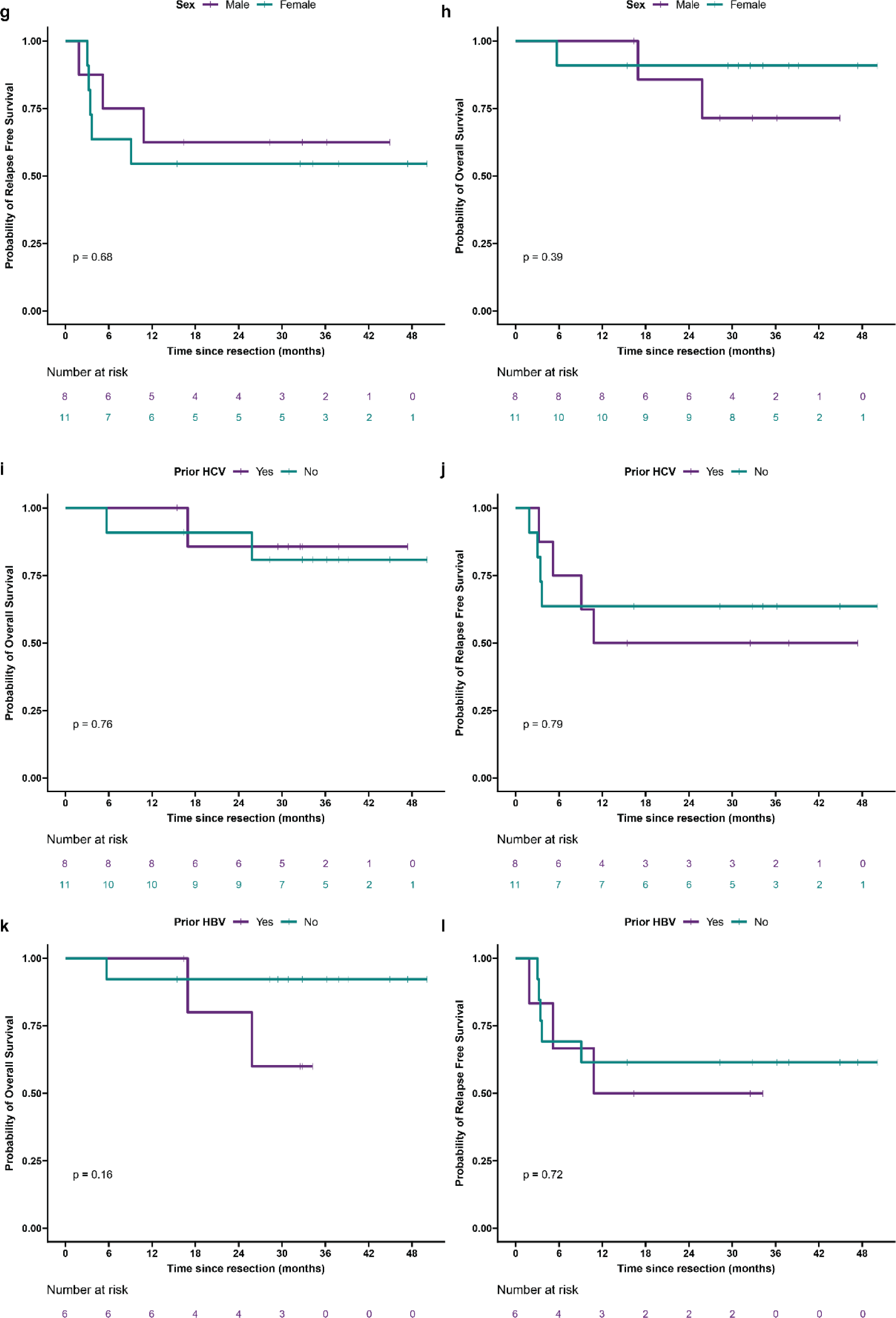
Relapse free survival and overall survival in HCC cohort treated with neoadjuvant ICB, according to clinical covariates. **a—l**, Kaplan-Meier curves showing relapse free survival and overall survival after surgical resection for HCC patients treated with neoadjuvant ICB (*n* = 19), according to total TLS density (**a** and **b**), peritumoral TLS density (**c** and **d**), pathologic response (**e** and **f**), sex (**g** and **h**), prior hepatitis C (HCV) infection (**i** and **j**), and prior hepatitis B (HBV) infection (**k** and **l**). Statistical significance was determined by log-rank test.

**Extended Data Fig. 3.**
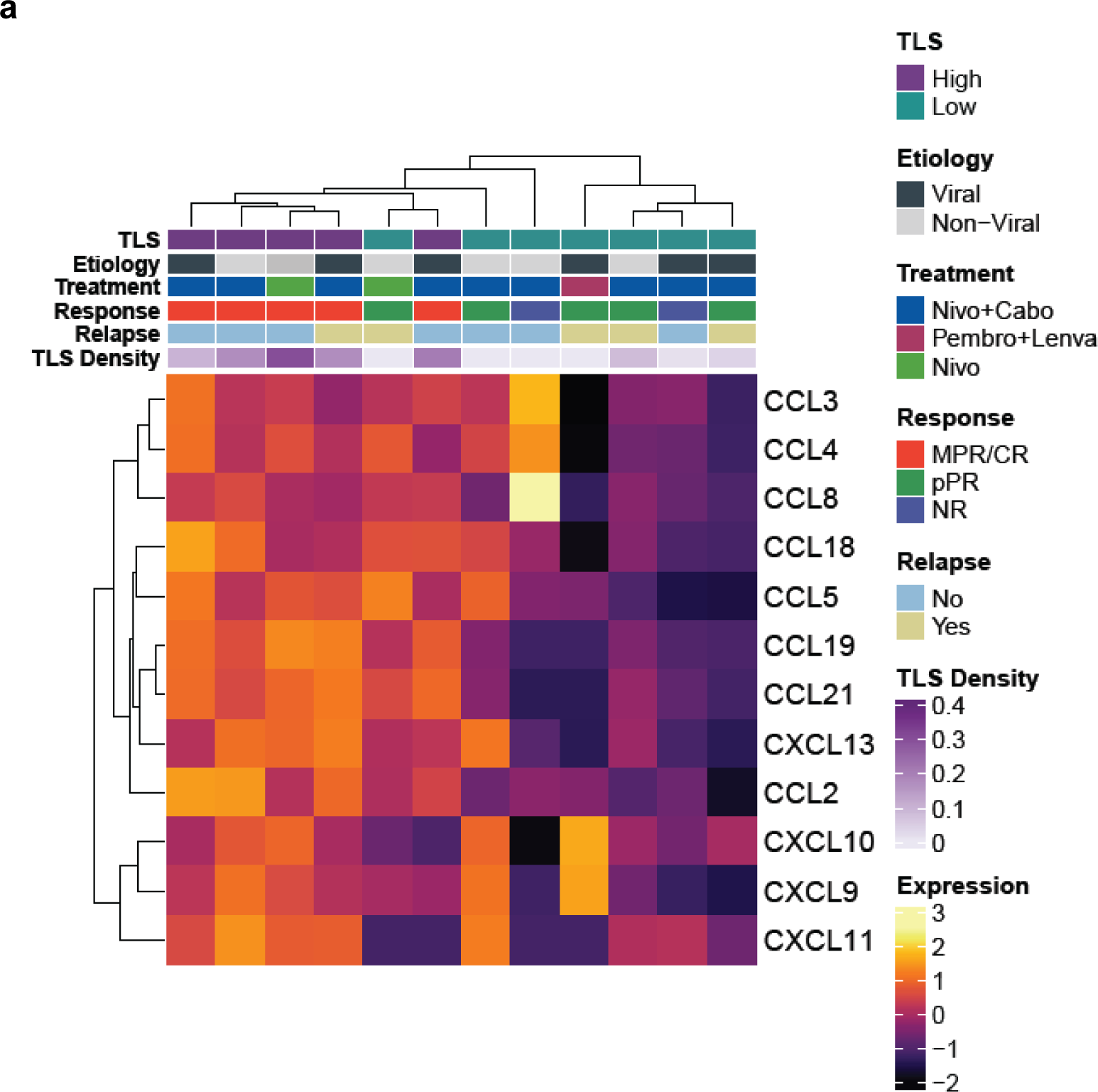
High TLS density after neoadjuvant ICB is associated with increased expression of the 12-chemokine TLS gene signature. a, Heatmap showing expression of the 12-chemokine gene signature in tumors with high TLS density (*n* = 5) and low TLS density (*n* = 7). Annotation rows indicate TLS group, HCC etiology, neoadjuvant treatment, pathologic response, relapse, and TLS density.

**Extended Data Fig. 4.**
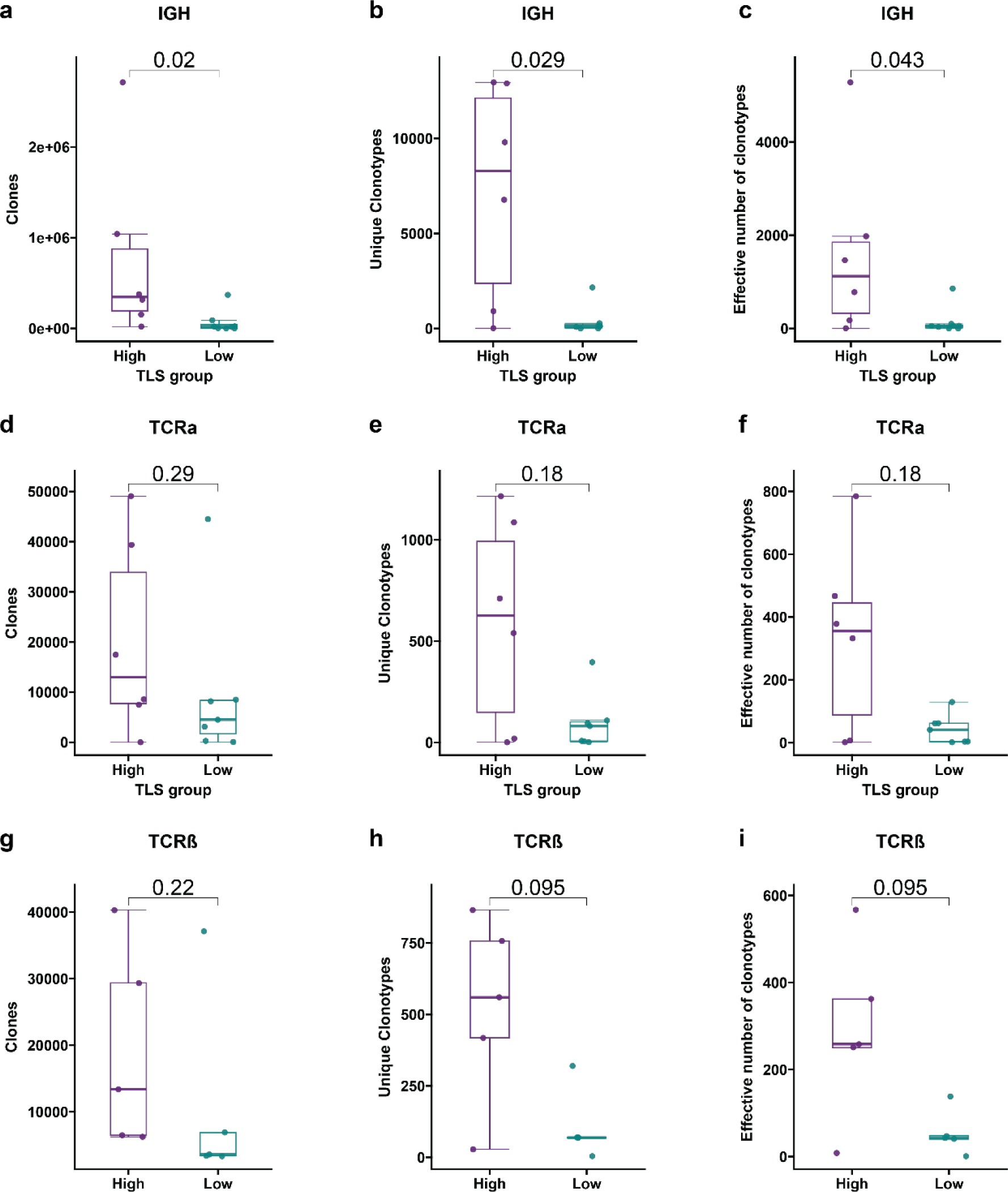
HCC tumors with high TLS density after neoadjuvant ICB have expanded T and B cell repertoires compared to tumors with low TLS density. Box-and-whisker plots showing the total clones, unique clonotypes, and effective number of clonotypes (i.e. true diversity index) for the immunoglobulin heavy chain (IGH) **(a-c),** TCRα (d-f), and TCRβ (**g-i**) repertoires of HCC tumors with high and low TLS density after neoadjuvant ICB. For each box-and-whisker plot, the horizontal bar indicates the median, the upper and lower limits of the boxes the interquartile range, and the ends of the whiskers 1.5 times the interquartile range. Statistical significance was determined by Wilcoxon rank sum test.

**Extended Data Fig. 5.**
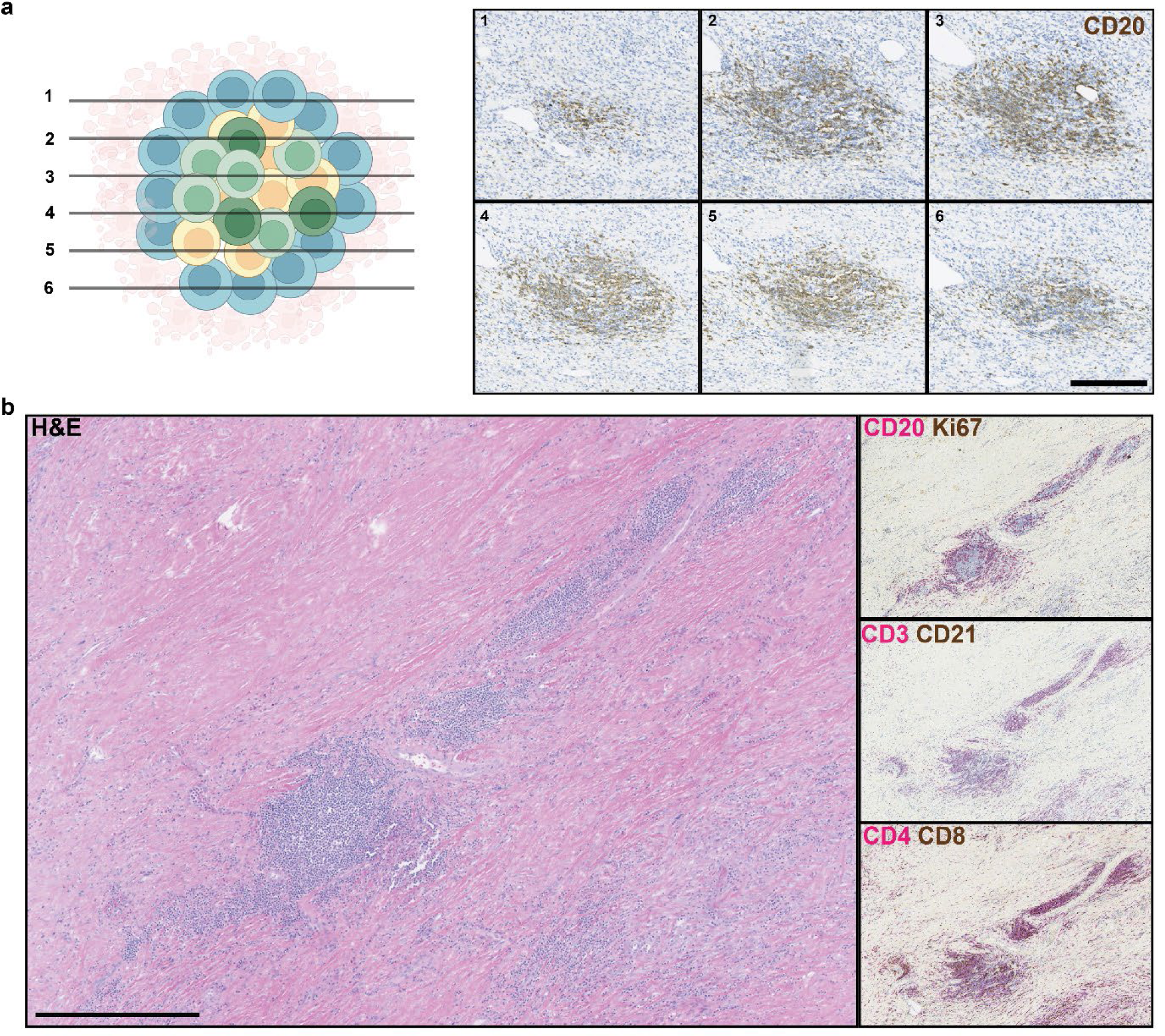
Involuted TLS in an HCC tumor with complete pathologic response after neoadjuvant ICB (OT7). **a**, Serial FFPE sections of an involuted TLS stained with anti-CD20 antibody (brown). Numbered images indicate the order in which the sections were cut from the tissue block. Scale bar, 250 μm. **b,** Representative images of multiple involuted TLS (red arrows) stained with hematoxylin and eosin (H&E), anti-CD20 (magenta) and anti-Ki67 (brown) (right middle), anti-CD3 (magenta) and anti-CD21 (brown) (middle right), and anti-CD4 and anti-CD8 (bottom right).

**Extended Data Fig. 6.**
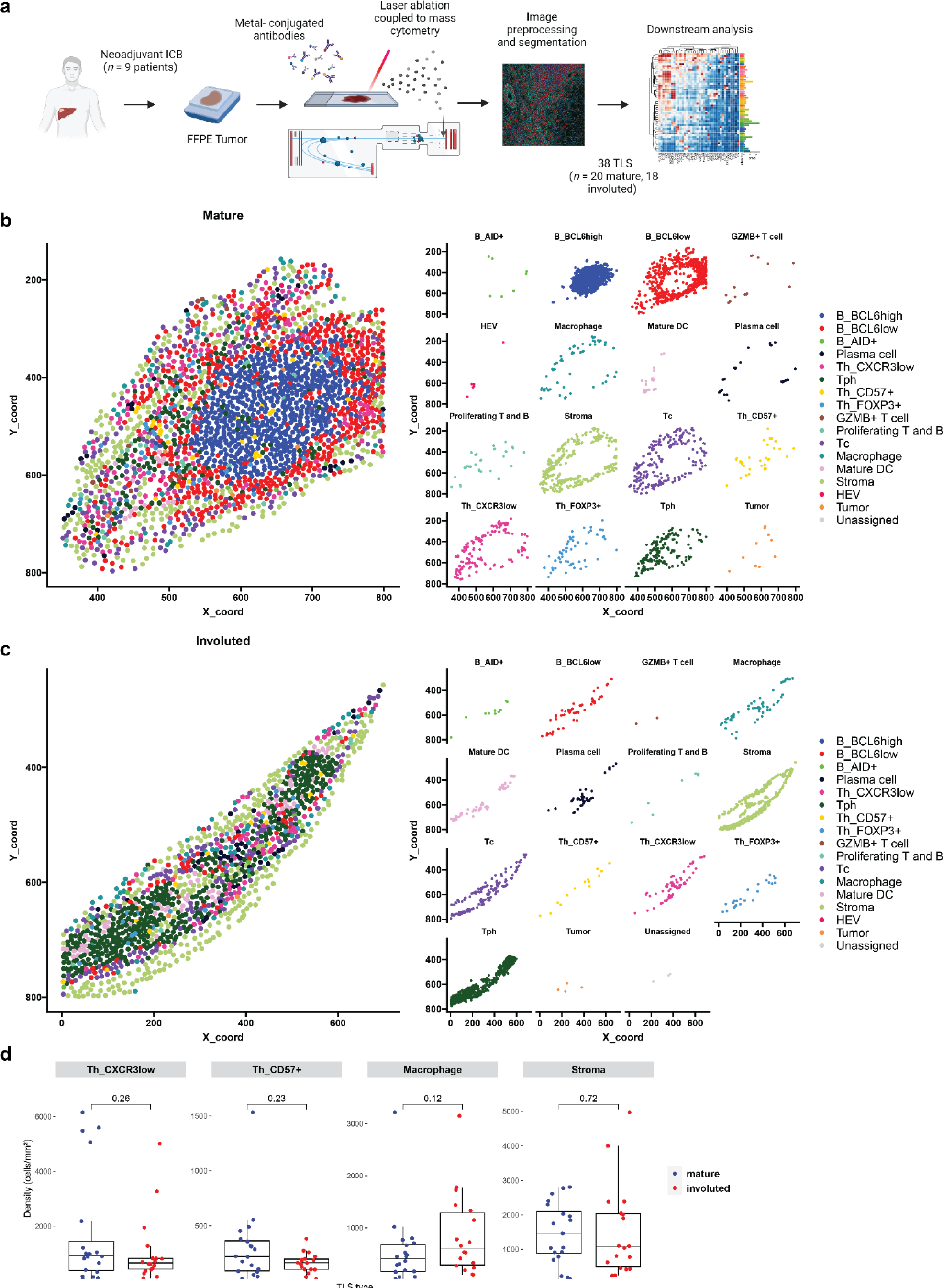
Characterization of divergent TLS morphologies in viable tumor and tumor regression bed by imaging mass cytometry. **a**, Imaging mass cytometry workflow. **b-c**, Dot plots showing representative mature (**b**) and involuted (**c**) TLS, colored according to cluster assignment of individual cells after cell segmentation. (**d**) Box-and-whisker plots showing cell cluster density in mature versus involuted TLS for CXCR3^low^ CD4 T cells, CD57^+^ CD4 T cells, Macrophages, and Stroma. For each box-and-whisker plot, the horizontal bar indicates the median, the upper and lower limits of the boxes the interquartile range, and the ends of the whiskers 1.5 times the interquartile range. Statistical significance was determined by pairwise two sample Wilcoxon test (**d**).

**Extended Data Fig. 7.**
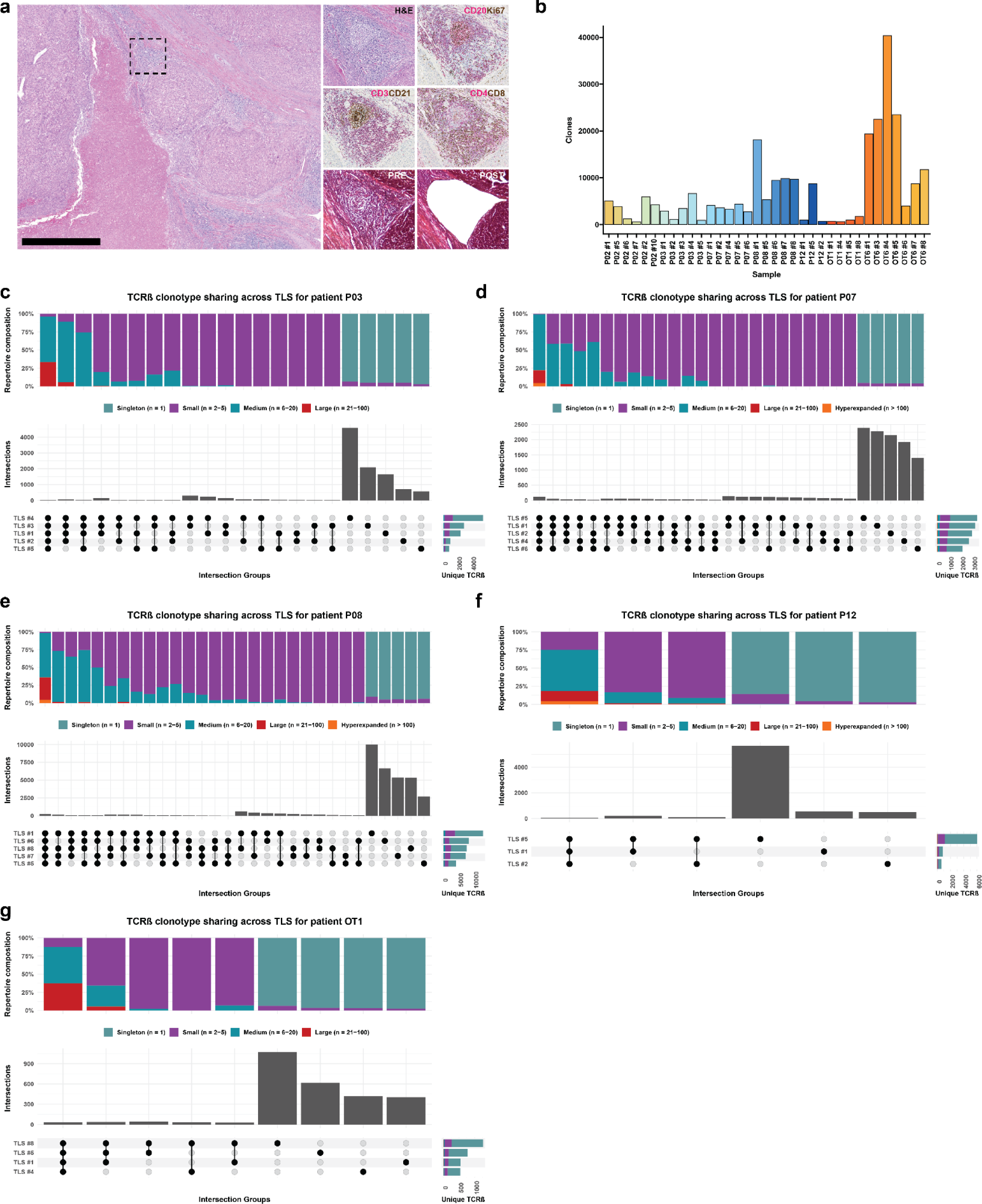
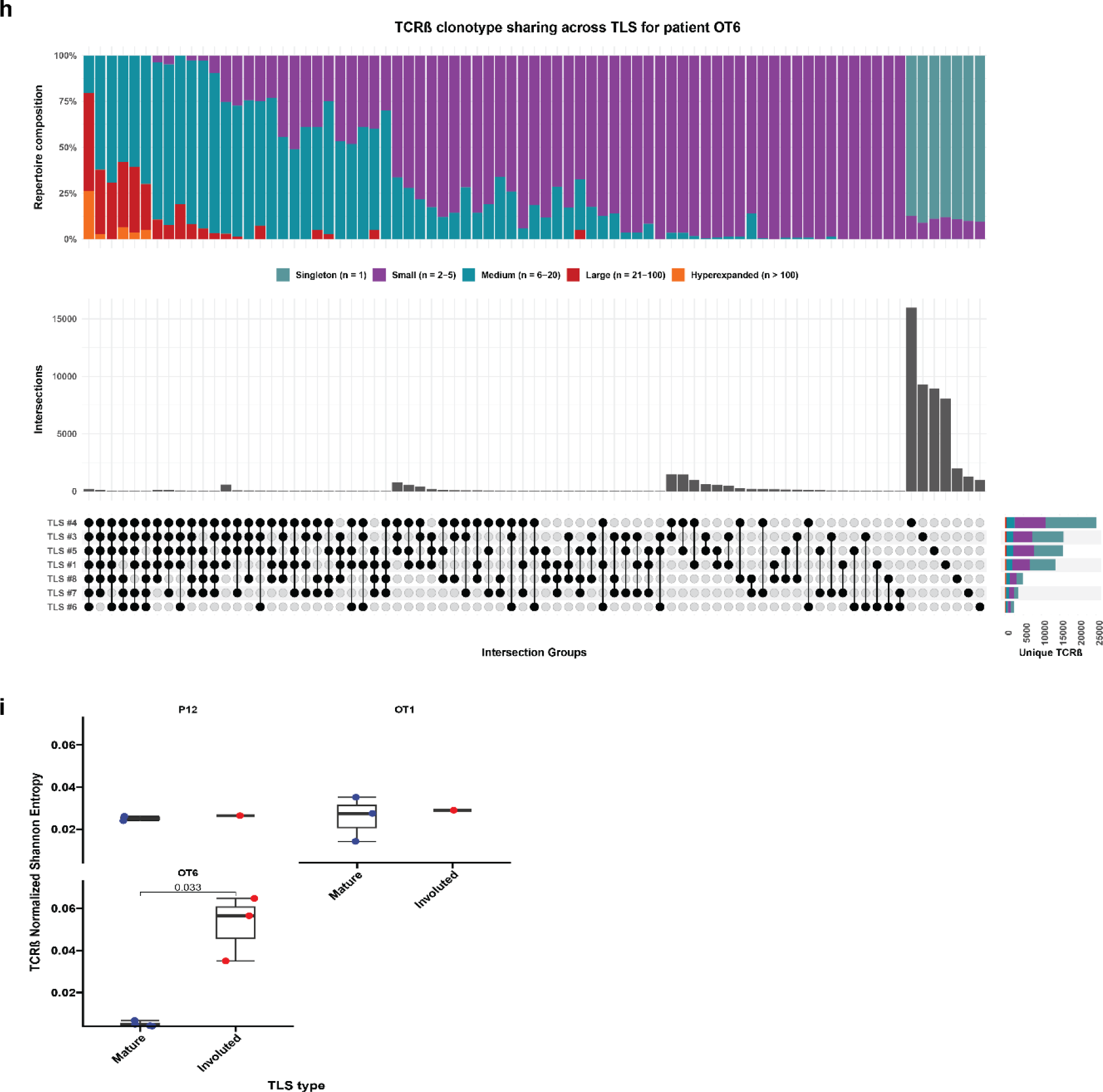
TCRβ repertoire features of microdissected TLS. **a**, Representative images showing method of identification and microdissection of individual TLS. Image on left shows HCC tumor stained with hematoxylin and eosin (H&E) at low magnification. Insets show higher magnification of staining with H&E, anti-CD20 (magenta) and anti-Ki67 (brown), anti-CD3 (magenta) and anti-CD21 (brown), anti-CD4 and anti-CD8 (bottom right), and corresponding pre- and post-microdissection images. Scale bar, 1mm. **b,** Barplot showing total clone count across all microdissected TLS. **c-h,** Representative upset plots showing overlap in TCRβ clonotypes across microdissected TLS from patients P03 (**c**), P07 (**d**), P08 (**e**), P12 (**f**), OT1 (**g**), and OT6 (**h**). For each upset plot, barplots in gray and row below indicate number of overlapping clonotypes between different combinations of TCRβ repertoires. Stacked barplots at top indicate repertoire composition of different groups of TCRβ and at bottom right indicate total number of unique TCRβ clonotypes identified in each TLS, colored according to clonal expansion. Intersections with fewer than 20 unique clonotypes are not shown. **i**, Dotplot showing TCRβ repertoire clonality (as determined by Normalized Shannon Entropy) for matched mature and involuted TLS. Statistical significance was determined by two-tailed t test (**i**).

**Extended Data Fig. 8.**
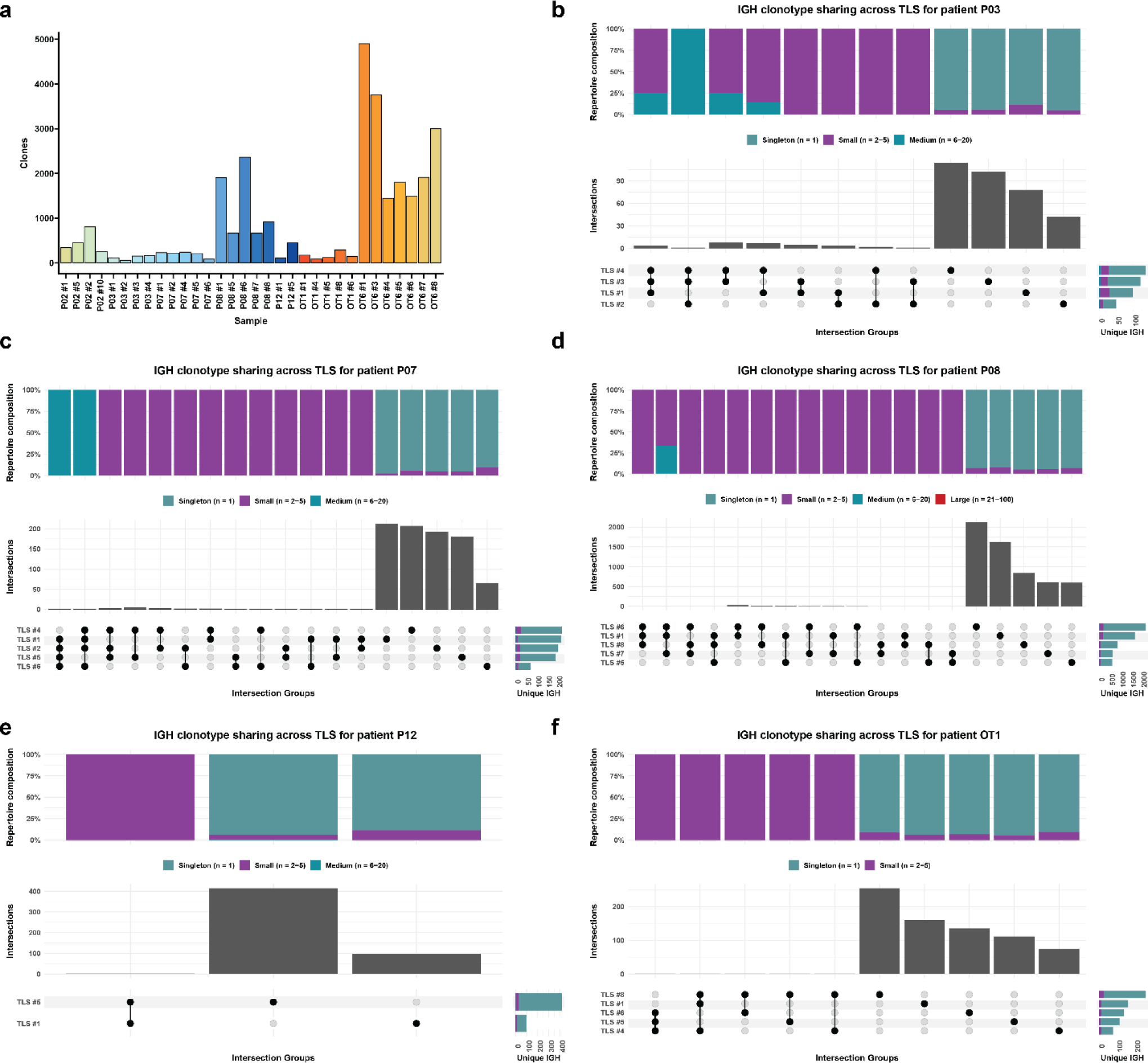
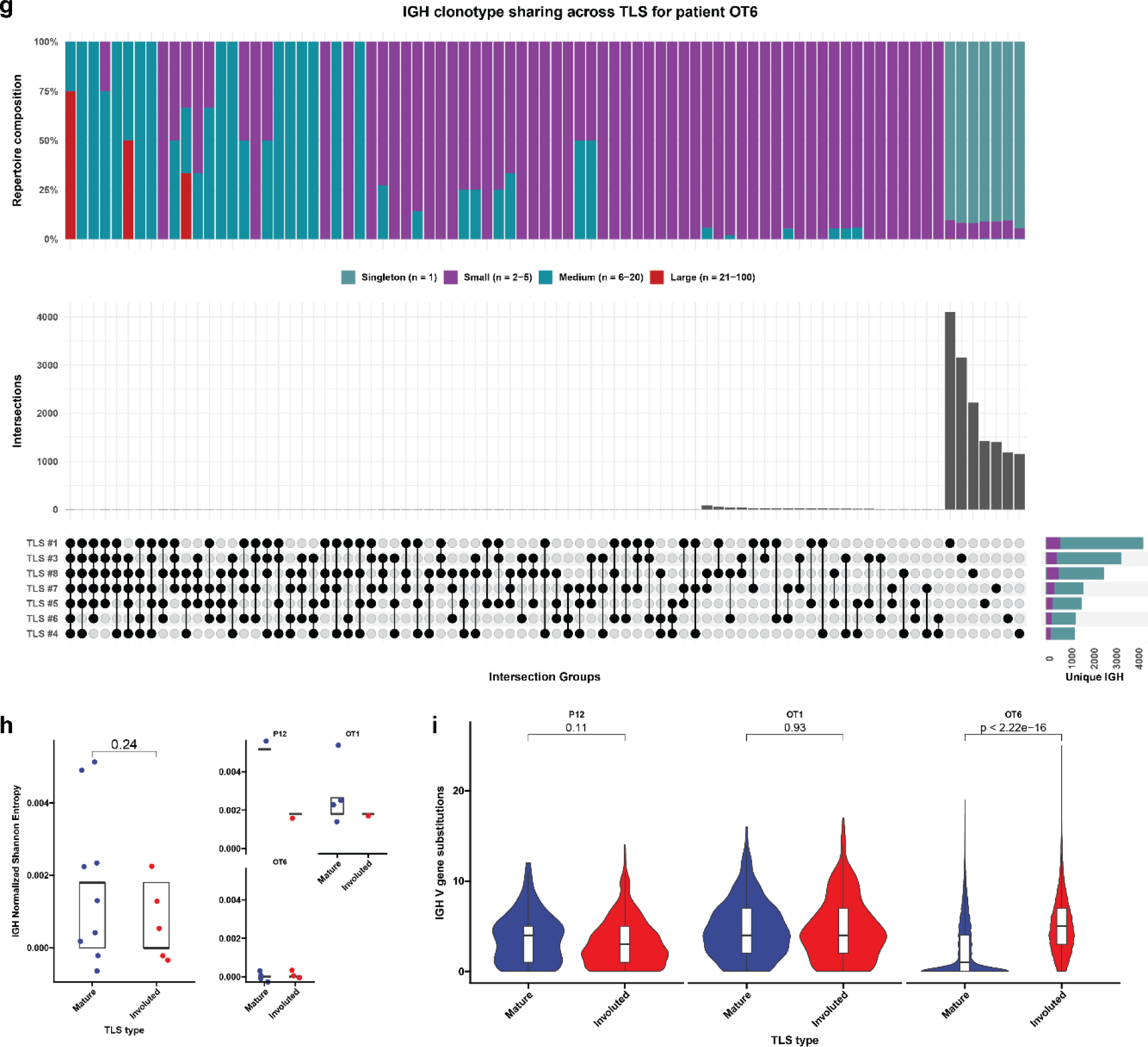
IGH repertoire features of microdissected TLS. **a**, Stacked barplot showing IGH repertoire composition across all TLS. **b**-**f,** Representative upset plots showing overlap in unique IGH clonotypes across microdissected TLS from patients P03 (**b**), P07 (**c**), P08 (**d**), P12 (**e**), OT1 (**f**), and OT7 (**g**). Bottom barplots and annotation row indicate number of overlapping clonotypes between different TLS repertoires. Top stacked barplots indicate clonal composition of overlapping (“public IGH”) and nonoverlapping (“Private IGH”). Bottom right stacked barplots indicate total number of unique IGH clonotypes identified at each TLS and overall clonal composition. **h,** Dotplot showing IGH repertoire clonality (as determined by Normalized Shannon Entropy) for microdissected TLS, according to TLS morphology. Statistical significance was determined by two-tailed t test (**h** and **i**).

**Extended Data Fig. 9.**
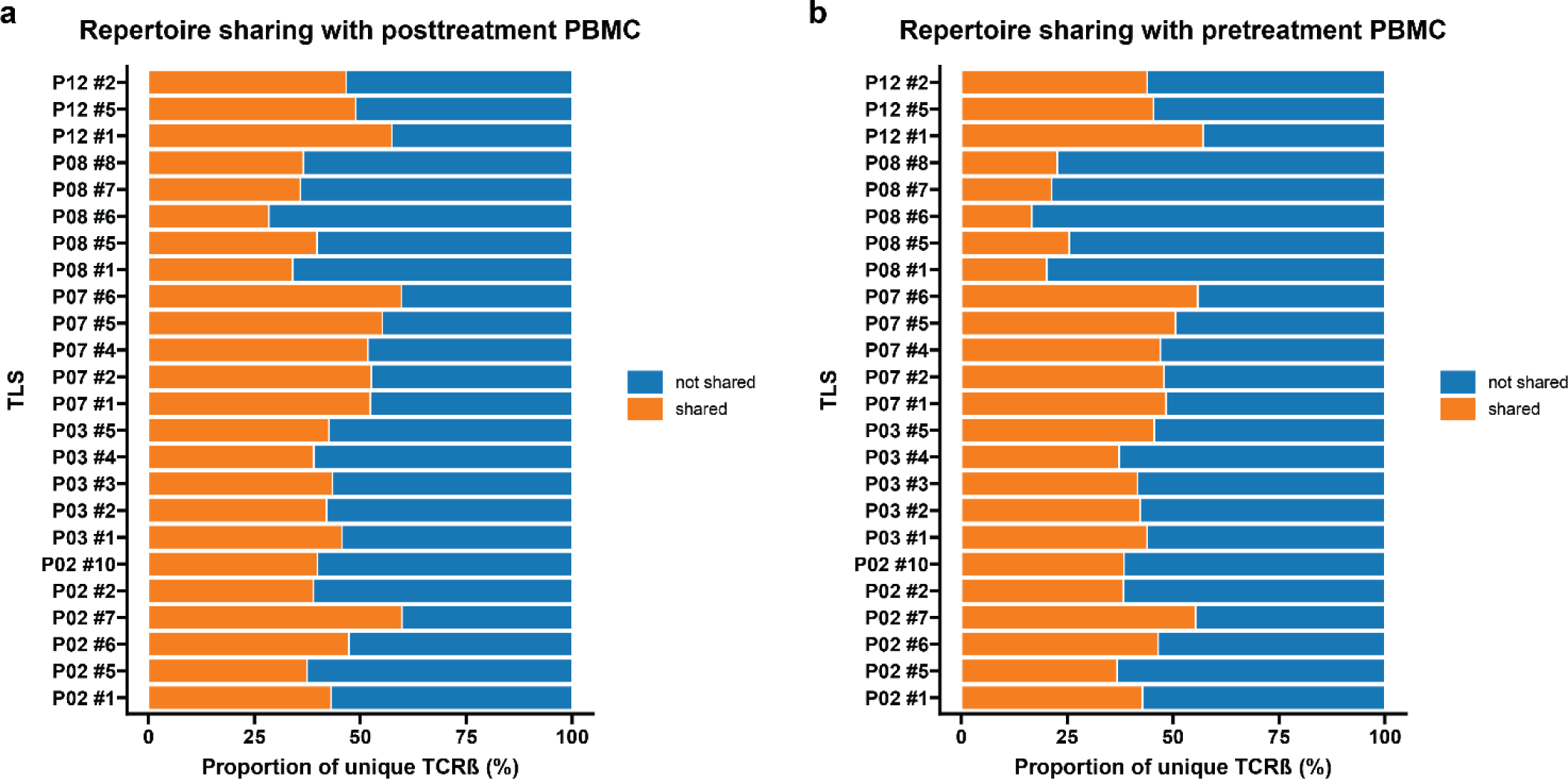
TLS display a high degree of T cell repertoire overlap with pre- and post-treatment peripheral blood. **a-b**, Barplots showing proportion of unique TCRβ clonotypes at each TLS that also identified in matched pre-treatment (**a**) and post-treatment (**b**) peripheral blood.

**Extended Data Fig. 10.**
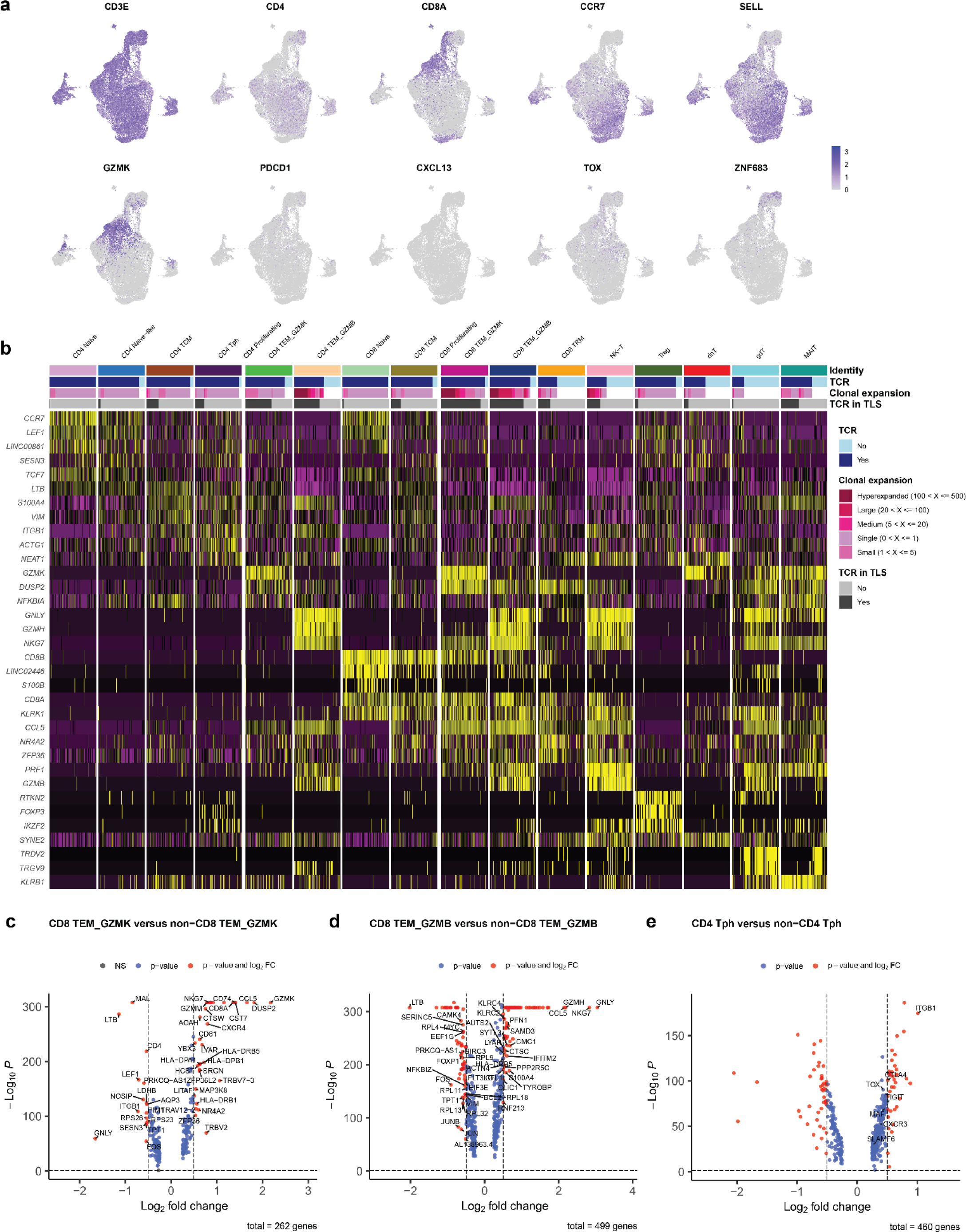
Single cell sequencing of post-treatment peripheral blood. a, UMAPs showing gene expression of *CD3E*, *CD4*, *CD8A*, *CCR7*, *SELL*, *GZMK*, *PDCD1*, *CXCL13*, *TOX*, and *ZNF683* across all single cells sequenced from post-treatment peripheral blood of 7 HCC patients treated with neoadjuvant ICB. **b,** Heatmap showing gene expression of the top 3 differentially expressed genes per cluster. Rows represent single genes and columns represent individual cells. Annotation bar indicates cluster identity, whether each cell had a sequenced TCR, the clonality of the TCR, and whether the TCR was identified in microdissected TLS from the same patient. Clusters were downsampled to 75 cells per cluster for visualization. **c-e,** Volcano plots showing differentially expressed genes in the CD8 TEM_GZMK (**b**),CD8 TEM_GZMB (**c**), and CD4 Tph **(d)** clusters compared to all other cells. Vertical dotted lines indicates a fold change of greater or less than 1.4 and horizontal line indicates a P value of 0.05. Labeled genes in **c** and **d** indicate genes with the highest differential expression. Labeled genes in **e** indicate genes known to be highly expressed in CD4 Tph.

**Extended Data Fig. 11.**
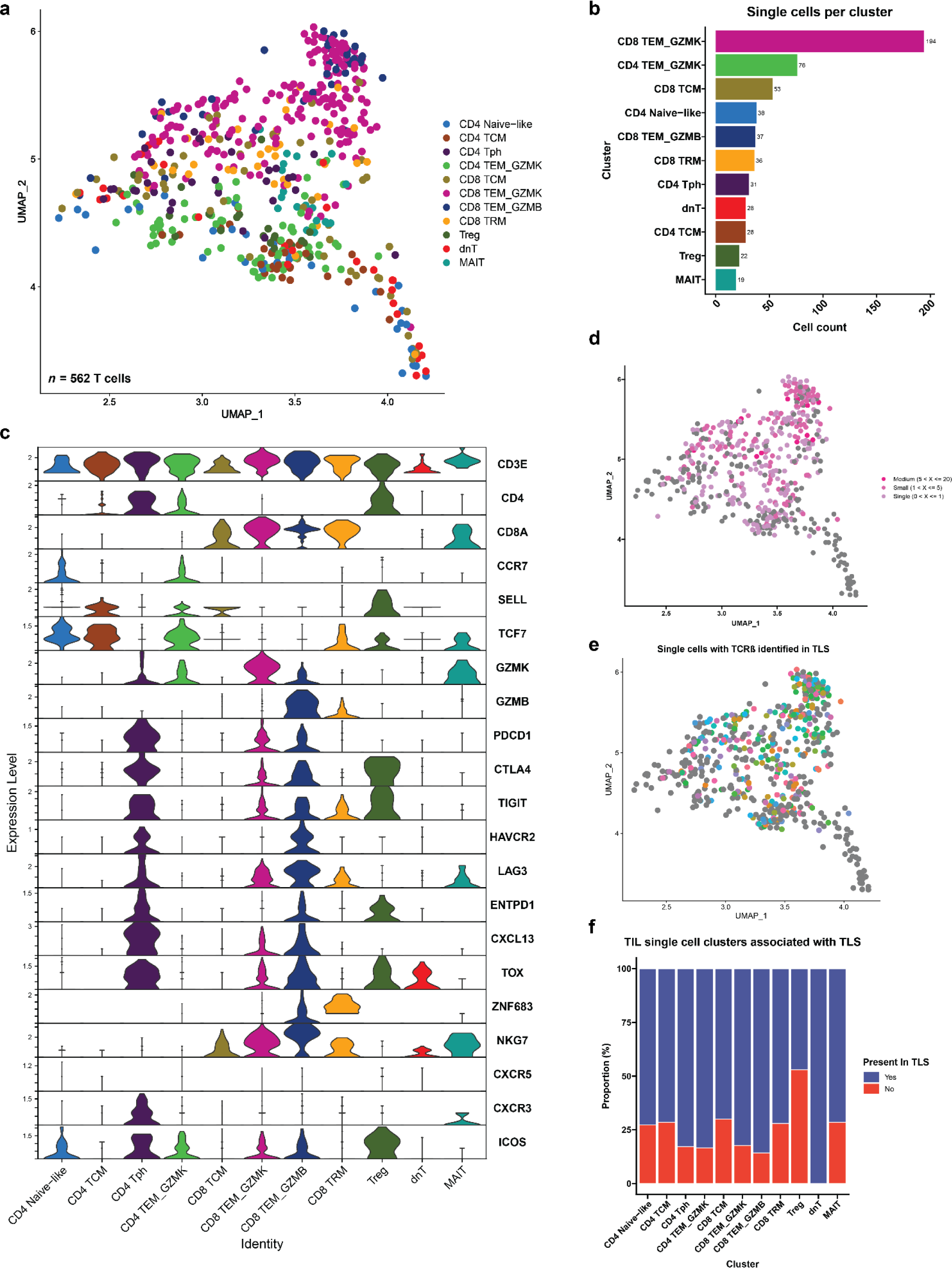
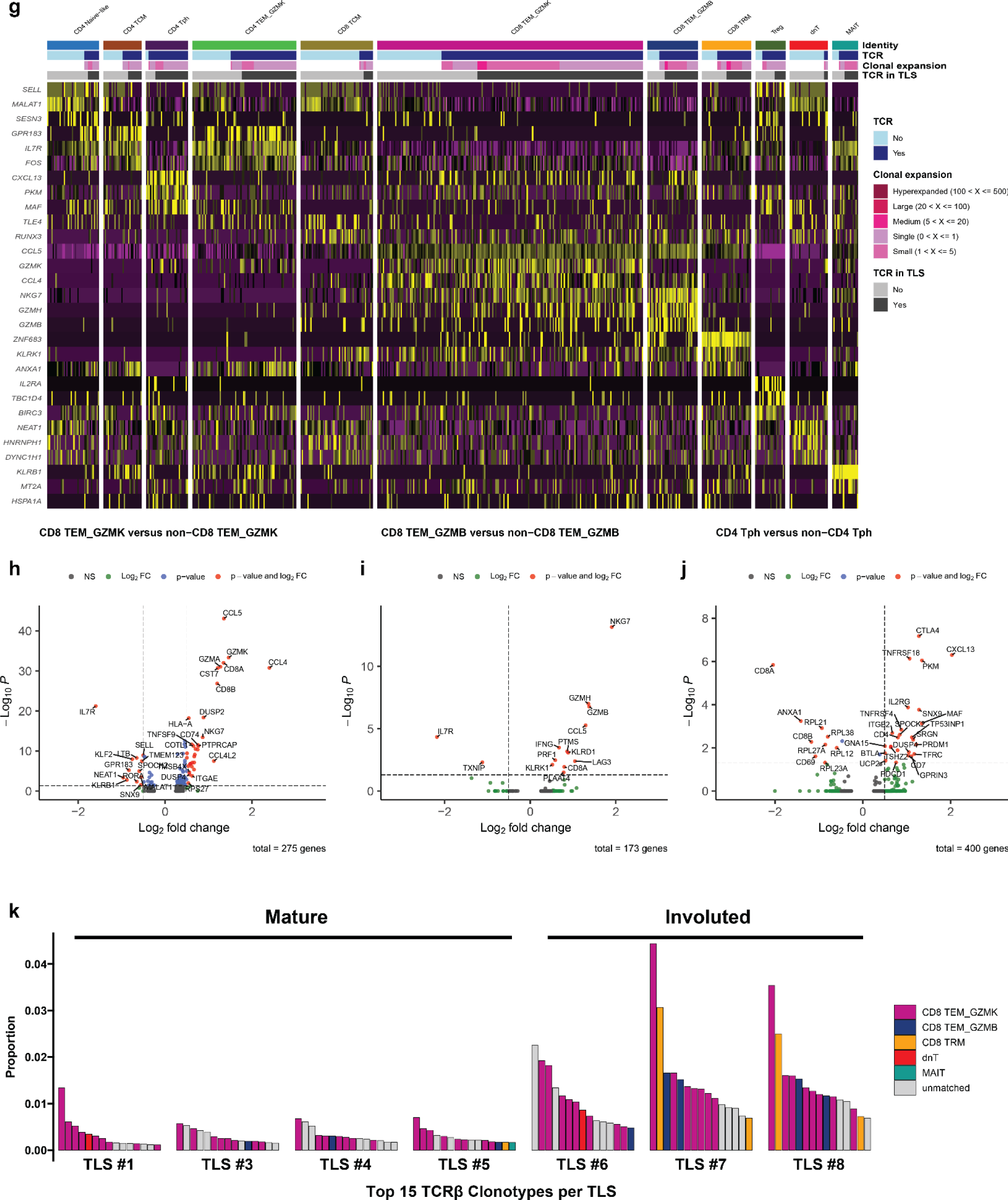
Single cell sequencing of post-treatment TIL from patient OT6. **a**, Uniform Manifold Approximation and Projection (UMAP) for 562 T cells identified by single cell RNA/TCR/BCR sequencing of CD3^+^CD19^+^ FACS-sorted tumor infiltrating lymphocytes. **b,** Barplot showing number of single cells per cluster. **c,** Violin plots showing expression of subset specific marker genes across clusters. **d-e,** UMAPs showing clonality of single cells with an associated T cell receptor sequence (**d**) and single cells with a TCRβ identified in microdissected TLS (**e**). **f,** Stacked barplot showing proportion of each single cell cluster identified in TLS. **g,** Heatmap showing gene expression of the top 3 differentially expressed genes per cluster. Rows represent single genes and columns represent individual cells. Annotation bar indicates cluster identity, whether each cell had a sequenced TCR, the clonality of the TCR, and whether the TCR was identified in microdissected TLS from the same patient. **h-j,** Volcano plots showing differentially expressed genes in the CD8 TEM_GZMK (**h**),CD8 TEM_GZMB (**i**), and CD4 Tph **(j)** clusters compared to all other cells. Vertical dotted lines indicates a fold change of greater or less than 1.4 and horizontal line indicates a P value of 0.05. **k,** Inferred transcriptional phenotype of the top 15 TCRβ clonotypes in mature and involuted TLS of patient OT6.

**Extended Data Fig. 12.**
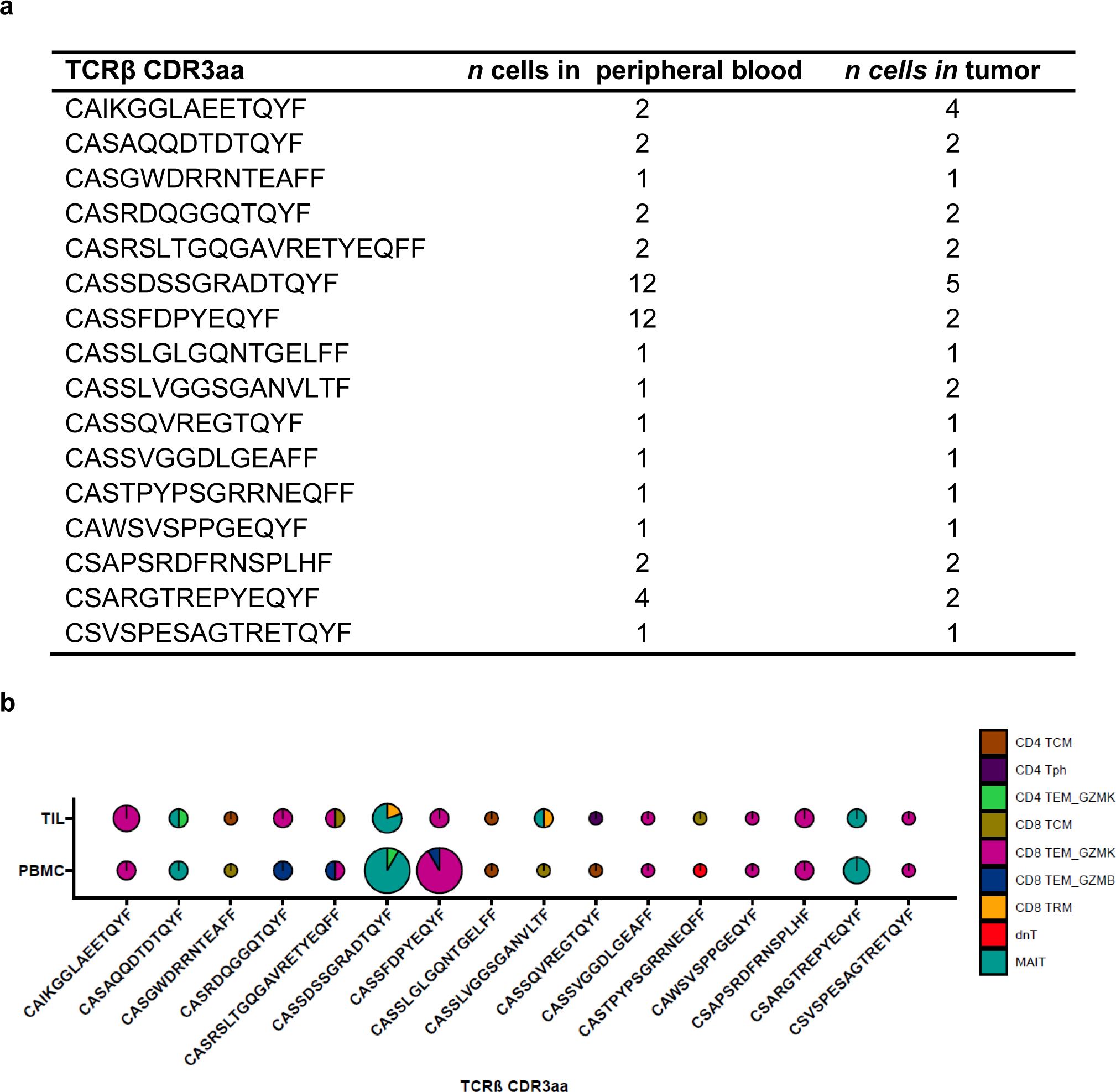
Cluster annotation of single cells with shared TCRβ in post-treatment peripheral blood and TIL (*n* = 16) from patient OT6. **a**, Shared TCRβ identified in both PBMC and TIL for patient OT6. Rows indicate different TCRβ clonotype and columns provide the complementarity determining region 3 (CDR3) amino acid sequence and number of cells with the TCRβ CDR3 amino acid sequence in peripheral blood and TIL, respectively. **b,** Single cell cluster identities of shared TCRβ according to unique CDR3 and compartment where the TCR was identified. Piecharts are colored according to the cluster identities of all cells with the same TCRβ. The radius of each piechart is proportional to the total number of cells in which each TCRβ was identified (square root of *n* cells divided by eight).

**Extended Data Table 1.** Clinical Characteristics of HCC cohort treated with neoadjuvant ICB. Characteristics of treated patients. Each row represents a single patient and columns indicate age at surgery, sex, HCC etiology, histologic grade, treatment regimen, pathologic response, and whether the patient suffered relapsed or death.

| Patient | Age at surgery | Sex | Etiology | Histologic grade | Treatment | Response | Relapse | Death |
| --- | --- | --- | --- | --- | --- | --- | --- | --- |
| P01 | 66 | M | HCV | Moderately differentiated | α-PD1 + TKI | NR | No | No |
| P02 | 72 | M | HCV | Moderately differentiated | α-PD1 + TKI | MPR | No | No |
| P03 | 76 | F | Unknown | Moderately differentiated | α-PD1 + TKI | MPR | No | No |
| P07 | 64 | F | HBV/HCV | Moderately differentiated | α-PD1 + TKI | pPR | Yes | Yes |
| P08 | 65 | M | HCV | Moderately differentiated | α-PD1 + TKI | MPR | Yes | No |
| P09 | 47 | M | HBV | Moderately differentiated | α-PD1 + TKI | CR | No | No |
| P10 | 41 | F | HBV | Moderately differentiated | α-PD1 + TKI | pPR | Yes | Yes |
| P11 | 56 | F | Unknown | Well differentiated | α-PD1 + TKI | pPR | No | No |
| P12 | 69 | M | HCV | Moderately differentiated | α-PD1 + TKI | MPR | No | No |
| P13 | 74 | F | Unknown | Poorly differentiated | α-PD1 + TKI | NR | No | No |
| P14 | 79 | M | Unknown | Moderately differentiated | α-PD1 + TKI | pPR | Yes | No |
| P15 | 49 | M | NASH | Moderately differentiated | α-PD1 + TKI | NR | Yes | Yes |
| OT1 | 68 | M | ETOH | Poorly differentiated | α-PD1 | MPR | No | No |
| OT2 | 71 | F | Unknown | Well differentiated | α-PD1 | pPR | No | No |
| OT3 | 63 | M | HCV | Moderately differentiated | α-PD1 + TKI | pPR | Yes | No |
| OT4 | 69 | F | HCV | Well differentiated | α-PD1 + TKI | pPR | Yes | No |
| OT5 | 62 | M | NASH | Moderately differentiated | α-PD1 | pPR | Yes | No |
| OT6 | 61 | M | HCV | Moderately differentiated | α-PD1 + α-CTLA4 + TKI | MPR | No | No |
| OT7 | 56 | F | HBV | Poorly differentiated | α-PD1 + α-CTLA4 | CR | No | No |

**Extended Data Table 2.** Clinical characteristics of untreated HCC cohort. Characteristics of untreated patients. Each row represents a single patient and columns indicate age at surgery, sex, HCC etiology, histologic grade, and treatment.

| Patient | Age at surgery | Sex | Etiology | Histologic grade | Treatment |
| --- | --- | --- | --- | --- | --- |
| C2 | 61 | M | HCV | Moderately differentiated | none |
| C3 | 60 | M | HBV/HCV | Well differentiated | none |
| C5 | 75 | M | HCV/NASH | Well differentiated | none |
| C6 | 63 | F | HCV | Moderately differentiated | none |
| C7 | 61 | M | HCV/ETOH | Well differentiated | none |
| C8 | 57 | M | NASH | Well differentiated | none |
| C9 | 65 | M | HBV/ETOH | Moderately differentiated | none |
| C10 | 66 | M | HCV | Well differentiated | none |
| C11 | 69 | F | HCV | Moderately differentiated | none |
| C12 | 71 | M | NASH | Moderately differentiated | none |
| C13 | 78 | M | Unknown | Moderately differentiated | none |
| C14 | 66 | M | Unknown | Moderately differentiated | none |
| C15 | 84 | F | NASH | Poorly differentiated | none |
| C16 | 49 | M | ETOH | Moderately differentiated | none |

**Extended Data Table 3.** Immune Related Pathologic Response Criteria Scoring for HCC tumors treated with neoadjuvant anti-PD-1.

| Feature, *n* (%) | MPR/CR (*n* = 8) | pPR/NR (*n* = 11) | P value† |
| --- | --- | --- | --- |
| **Tumor** |  |  |  |
| Fibrosis |  |  |  |
| Immature fibrosis | 8 (100%) | 7 (63.6%) | 0.103 |
| Mature fibrosis | 8 (100%) | 7 (63.6%) | 0.103 |
| Neovascularization | 1 (12.5%) | 0 (0%) | 0.421 |
| Cholesterol clefts | 4 (50.0%) | 2 (18.2%) | 0.319 |
| Granulomas | 3 (37.5%) | 2 (18.2%) | 0.603 |
| Foamy histiocytes | 5 (62.5%) | 4 (36.4%) | 0.37 |
| Giant cells | 1 (12.5%) | 2 (18.2%) | 1 |
| Hemosiderin macrophages | 7 (87.5%) | 5 (45.5%) | 0.147 |
| Calcification | 1 (12.5%) | 1 (9.1%) | 1 |
| Lymphoid aggregates | 8 (100%) | 8 (72.7%) | 0.228 |
| Tertiary lymphoid structures | 6 (75.0%) | 2 (18.2%) | 0.0237 |
| Dense plasma cells | 2 (25.0%) | 1 (9.1%) | 0.546 |
| **Peritumor** |  |  |  |
| Lymphoid aggregates | 7 (87.5%) | 8 (72.7%) | 0.603 |
| Tertiary lymphoid structures | 4 (50.0%) | 3 (27.3%) | 0.377 |
| Dense plasma cells | 1 (12.5%) | 0 (0%) | 0.421 |

† Fisher's Exact test

**Extended Data Table 4.** Bayesian Information Criteria results for predicting relapse and death following surgical resection in HCC treated with neoadjuvant ICB. Rows represent different BIC calculations and columns indicate outcomes and variables evaluated and calculated BIC.

| Outcome | Variable | BIC |
| --- | --- | --- |
| Relapse | Total TLS density | 43.04 |
| Relapse | Peritumoral TLS density | 45.4 |
| Relapse | Intratumoral TLS | 37.61 |
| Relapse | Pathologic Response | 40.18 |
| Relapse | Sex | 45.58 |
| Relapse | Prior HCV | 45.68 |
| Relapse | Prior HBV | 45.63 |
| Death | Total TLS density | 15.99 |
| Death | Peritumoral TLS density | 15.63 |
| Death | Intratumoral TLS | 15.99 |
| Death | Pathologic Response | 14.89 |
| Death | Sex | 17.2 |
| Death | Prior HCV | 17.85 |
| Death | Prior HBV | 16.22 |

**Extended Data Table 5 | Differentially expressed genes in TLS high and TLS low tumors.** Each row represents a single gene, and columns provide mean of normalized counts for all samples, log2 fold change in mRNA expression, Wald statistic, Wald test p-value, and Benjamini Hochberg adjusted p-values.

**Extended Data Table 6 | Gene Set Enrichment Results showing enriched pathways in TLS high tumors compared to TLS low tumors.** Each tab corresponds to a different gene set in the human MSigDB. Each row represents a single pathway, and columns provide the name of the pathway, enrichment p-values, a Benjamini Hochberg adjusted p-value, the expected error for the standard deviation of the P-value logarithm, enrichment score, enrichment score normalized to mean enrichment of random samples of the same size, size of the pathway after removing genes not present, and a vector with indexes of leading-edge genes that drive the enrichment.

**Extended Data Table 7.**
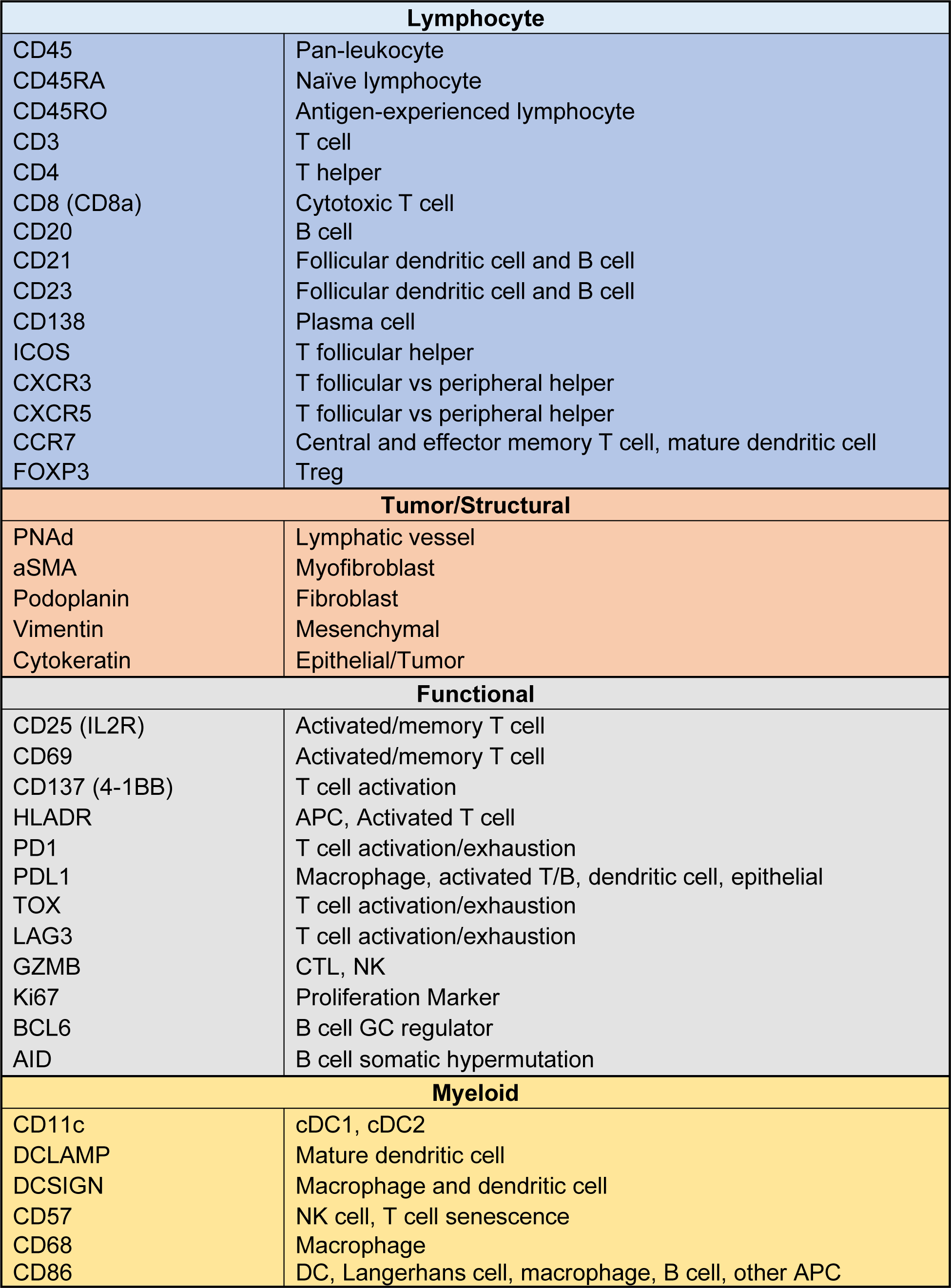
Summary of antibodies selected for imaging mass cytometry.

**Extended Data Table 8.** Summary of metal-conjugated antibodies used for imaging mass cytometry. Rows indicates a different stain, and columns indicate the metal, antigen, clone, dilution factor, source, and whether the antibody was custom-conjugated for the current study.

| Mass | Metal | Antigen | Clone | Dilution Factor | Source | Custom |
| --- | --- | --- | --- | --- | --- | --- |
| 89 | Y | CD45 | D9M8I | 125 | Cell Signaling Technology® | X |
| 96-104 | Ru | Counterstain |  |  | Electron Microscopy Sciences |  |
| 113 | In | PNAd | MECA-79 | 250 | Biolegend® | X |
| 115 | In | AID | mAID-2 | 125 | eBioscience™ | X |
| 141 | Pr | αSMA | 1A4 | 500 | Standard BioTools™ |  |
| 142 | Nd | Podoplanin | D2-40 | 125 | Biolegend® | X |
| 143 | Nd | Vimentin | D21H3 | 500 | Standard BioTools™ |  |
| 144 | Nd | CD11c | EP1347Y | 250 | Abcam | X |
| 145 | Nd | CD45RO | UCHL1 | 250 | Biolegend® | X |
| 146 | Nd | CXCR3 | IC6 | 100 | AbboMax | X |
| 147 | Sm | CD69 | EPR21814 | 250 | Abcam | X |
| 148 | Nd | Pan-Keratin | C11 | 125 | Standard BioTools™ |  |
| 149 | Sm | CD25 (IL2R) | SP176 | 250 | Abcam | X |
| 150 | Nd | PD-L1 | E1L3N | 125 | Cell Signaling Technology® | X |
| 151 | Eu | CXCR5 | 51505 | 250 | Novus Biologicals | X |
| 152 | Sm | DC-LAMP | 1010E1.01 | 125 | Novus Biologicals | X |
| 153 | Eu | Tox/Tox2 | E6I3Q | 250 | Cell Signaling Technology® | X |
| 154 | Sm | CD57 | HNK-1 | 250 | Cell Signaling Technology® | X |
| 155 | Gd | Foxp3 | PCH101 | 100 | Standard BioTools™ |  |
| 156 | Gd | CD4 | EPR6855 | 125 | Standard BioTools™ |  |
| 158 | Gd | ICOS | D1K2T™ | 250 | Cell Signaling Technology® | X |
| 159 | Tb | CD68 | KP1 | 100 | Standard BioTools™ |  |
| 160 | Gd | Syndecan 1 (CD138) | IHC138 | 125 | Cell Signaling Technology® | X |
| 161 | Dy | CD20 | H1 | 125 | Standard BioTools™ |  |
| 162 | Dy | CD8α | C8/144B | 250 | Standard BioTools™ |  |
| 163 | Dy | CD21 | Bu32 | 250 | Biolegend® | X |
| 164 | Dy | BCL-6 | K112-91 | 250 | BD Pharmingen™ | X |
| 165 | Ho | PD1 | EPR4877(2) | 250 | Abcam | X |
| 166 | Er | CD45RA | HI100 | 250 | Standard BioTools™ |  |
| 167 | Er | Granzyme B | D6E9W | 125 | Cell Signaling Technology® | X |
| 168 | Er | Ki-67 | B56 | 250 | Standard BioTools™ |  |
| 169 | Tm | CD23 | MRQ-57 | 125 | Cell Marque™ | X |
| 170 | Er | CD3ε | Polyclonal, C-terminal | 125 | Standard BioTools™ |  |
| 171 | Yb | Lag3 | 17B4 | 125 | GeneTex | X |
| 172 | Yb | CD137 (4-1BB) | D2Z4Y | 250 | Cell Signaling Technology® | X |
| 173 | Yb | DC-SIGN | DCN46 | 62.5 | lonpath | X |
| 174 | Yb | HLA-DR | LN3 | 250 | Standard BioTools™ |  |
| 175 | Lu | CD86 | E2G8P | 125 | Cell Signaling Technology® | X |
| 176 | Yb | CCR7 | EPR23192-57 | 250 | Cell Signaling Technology® | X |
| 191 | Ir | DNA 1 |  |  | Standard BioTools™ |  |
| 193 | Ir | DNA 2 |  |  | Standard BioTools™ |  |
| 195 | Pt | Plasma Membrane 2 | 1A36 | 250 | Standard BioTools™ |  |
| 196 | Pt | Plasma Membrane 3 | 1A37 | 250 | Standard BioTools™ |  |
| 198 | Pt | Plasma Membrane 4 | 1A38 | 250 | Standard BioTools™ |  |

**Extended Data Table 9.**
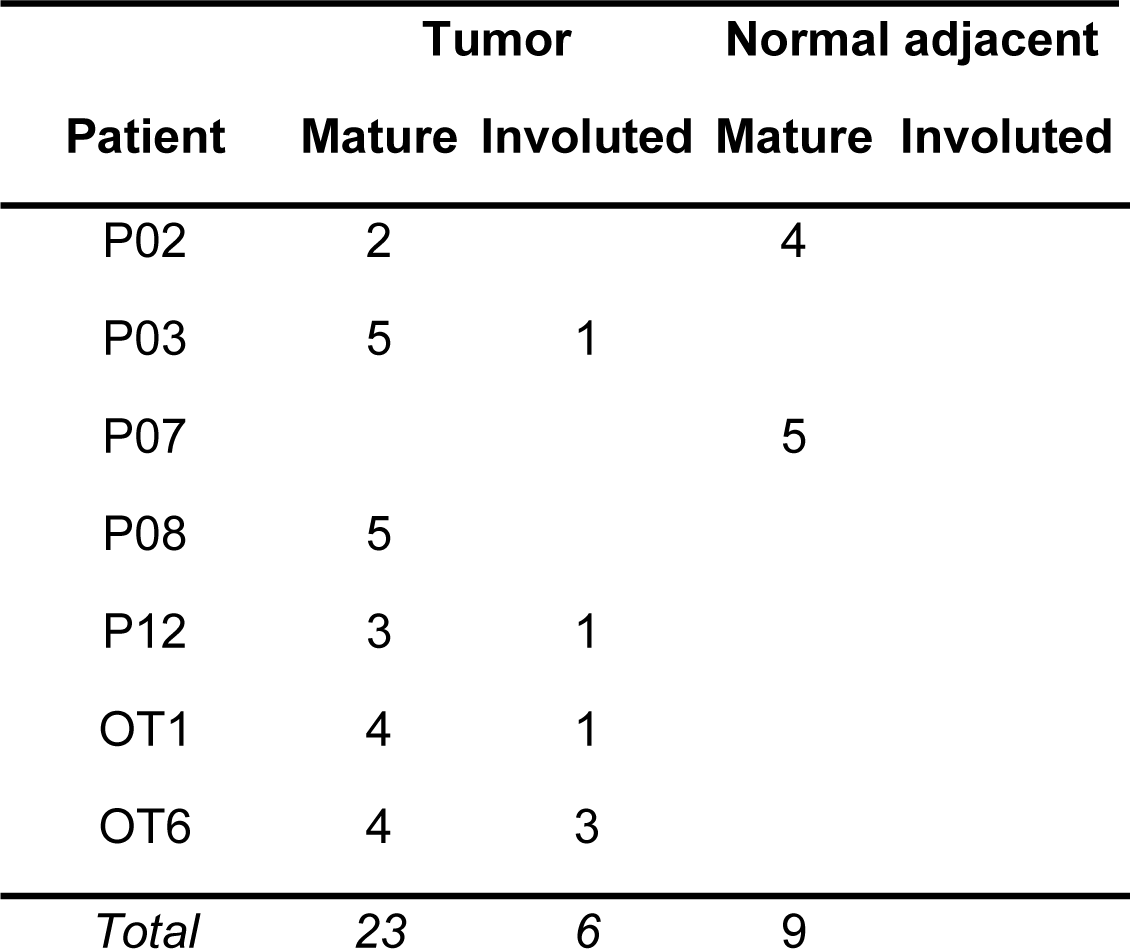
Characteristics of microdissected TLS. Each row indicates a different patient with HCC treated with neoadjuvant immunotherapy. Columns indicate number of TLS microdissected per patient according to location (tumor or normal adjacent) and morphology (mature or involuted). Sample number shown differs from the final number of TCRβ and IGH repertoires analyzed due to filtering to remove TCRβ repertoires with fewer than 500 clones and IGH repertoires with fewer than 50 clones.

| Patient | Tumor |  | Normal adjacent |  |
| --- | --- | --- | --- | --- |
|  | Mature | Involuted | Mature | Involuted |
| P02 | 2 |  | 4 |  |
| P03 | 5 | 1 |  |  |
| P07 |  |  | 5 |  |
| P08 | 5 |  |  |  |
| P12 | 3 | 1 |  |  |
| OT1 | 4 | 1 |  |  |
| OT6 | 4 | 3 |  |  |
| *Total* | 23 | 6 | 9 |  |

**Extended Data Table 10.**
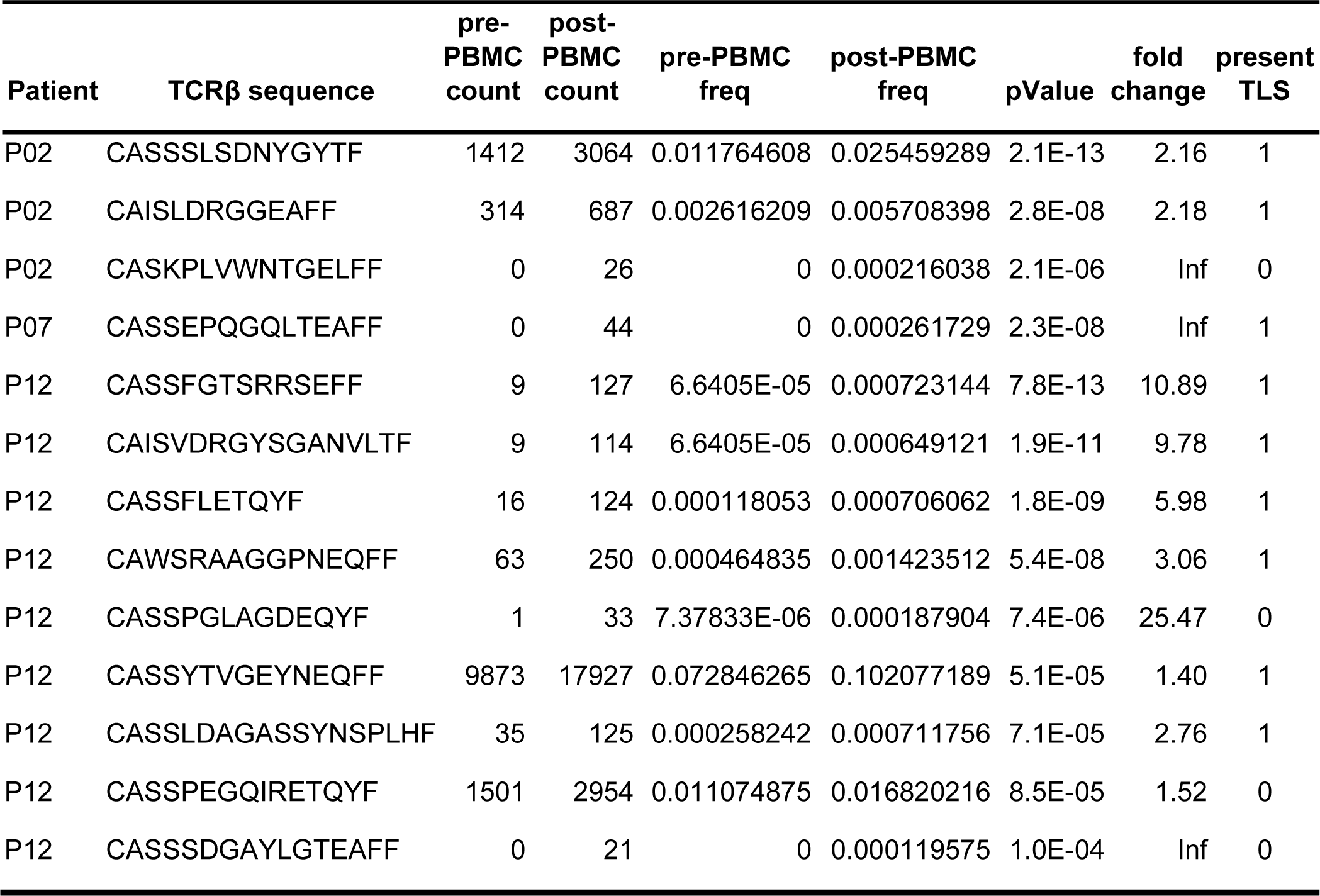
Expanded TCRβ clones in peripheral blood after neoadjuvant ICB.

**Extended Data Table 11.** TCR repertoire characteristics of peripheral blood and TIL single cell RNA/TCR-seq data.

| Cluster | Cells, *n* | Cells with TCR, *n* | Cells with expanded TCR, *n* (%) | Cells with TCRβ in TLS, *n* (%) |
| --- | --- | --- | --- | --- |
| CD4 Naive | 615 | 590 | 47 (8) | 16 (2.7) |
| CD8 Naive | 848 | 807 | 88 (10.9) | 11 (1.4) |
| MAIT | 315 | 217 | 108 (49.8) | 129 (59.4) |
| CD4 Naive-like | 8580 | 7224 | 806 (11.2) | 604 (8.4) |
| CD4 TCM | 2677 | 2531 | 464 (18.3) | 767 (30.3) |
| CD4 Tph | 1088 | 988 | 113 (11.4) | 188 (19) |
| CD4 TEM_GZMK | 1271 | 1086 | 384 (35.4) | 702 (64.6) |
| CD4 TEM_GZMB | 942 | 698 | 660 (94.6) | 601 (86.1) |
| CD8 TCM | 892 | 785 | 164 (20.9) | 202 (25.7) |
| CD8 TEM_GZMK | 1906 | 1676 | 1147 (68.4) | 1390 (82.9) |
| CD8 TEM_GZMB | 2551 | 1963 | 1682 (85.7) | 1569 (79.9) |
| CD8 TRM | 212 | 91 | 41 (45.1) | 52 (57.1) |
| NK-T | 75 | 32 | 25 (78.1) | 22 (68.8) |
| Treg | 801 | 712 | 106 (14.9) | 70 (9.8) |
| dnT | 267 | 116 | 23 (19.8) | 23 (19.8) |
| gdT | 132 | 30 | 7 (23.3) | 3 (10) |
| Total | 23172 | 19546 | 5865 (30) | 6349 (32.5) |

**TIL**
| Cluster | Cells, *n* | Cells with TCR, *n* | Cells with expanded TCRs, *n* (%) | Cells with TCRβ in TLS, *n* (%) |
| --- | --- | --- | --- | --- |
| CD4 Naive-like | 38 | 11 | 3 (27.3) | 8 (72.7) |
| CD4 TCM | 28 | 14 | 1 (7.1) | 10 (71.4) |
| CD4 Tph | 31 | 29 | 7 (24.1) | 24 (82.8) |
| CD4 TEM_GZMK | 76 | 48 | 10 (20.8) | 40 (83.3) |
| CD8 TCM | 53 | 10 | 3 (30) | 7 (70) |
| CD8 TEM_GZMK | 194 | 147 | 68 (46.3) | 121 (82.3) |
| CD8 TEM_GZMB | 37 | 28 | 16 (57.1) | 24 (85.7) |
| CD8 TRM | 36 | 25 | 14 (56) | 18 (72) |
| Treg | 22 | 17 | 3 (17.6) | 8 (47.1) |
| dnT | 28 | 3 | 1 (33.3) | 3 (100) |
| MAIT | 19 | 14 | 10 (71.4) | 10 (71.4) |
| Total | 562 | 346 | 136 (39.3) | 273 (78.9) |

**Extended Data Table 12.** Association of peripheral blood and TIL single cell clusters with clonal expansion.°†.

| Cluster | P value | OR | 95%_CI_Lower | 95%_CI_Upper |
| --- | --- | --- | --- | --- |
| CD4 TEM_GZMB | < 0.001 | 31.48 | 25.72 | 38.83 |
| CD8 TEM_GZMB | < 0.001 | 17.99 | 16.17 | 20.04 |
| NK-T | < 0.001 | 10.27 | 4.73 | 23.62 |
| Treg | < 0.001 | 9.02 | 5.42 | 16.23 |
| CD8 TEM_GZMK | < 0.001 | 6.27 | 5.63 | 6.97 |
| MAIT | 0.398 | 0.82 | 0.54 | 1.22 |
| CD8 TRM | 0.389 | 0.73 | 0.35 | 1.39 |
| gdT | 0.614 | 0.59 | 0.12 | 1.93 |
| CD4 TEM_GZMK | < 0.001 | 0.4 | 0.32 | 0.51 |
| dnT | 0.002 | 0.34 | 0.13 | 0.73 |
| CD8 TCM | < 0.001 | 0.28 | 0.2 | 0.38 |
| CD4 TCM | < 0.001 | 0.13 | 0.1 | 0.17 |
| CD8 Naive | < 0.001 | 0.08 | 0.04 | 0.14 |
| CD4 Naive | < 0.001 | 0.05 | 0.02 | 0.12 |
| CD4 Naive-like | < 0.001 | 0.04 | 0.03 | 0.05 |
| CD4 Tph | < 0.001 | 0.04 | 0.01 | 0.07 |

**TIL**
| Cluster | P value | OR | 95%_CI_Lower | 95%_CI_Upper |
| --- | --- | --- | --- | --- |
| Treg | 0.03 | Inf | 1.2 | Inf |
| CD8 TEM_GZMB | 0.003 | 3.49 | 1.45 | 8.32 |
| CD8 TRM | 0.056 | 2.25 | 0.88 | 5.49 |
| CD8 TEM_GZMK | 0.013 | 1.94 | 1.12 | 3.36 |
| MAIT | 0.518 | 1.44 | 0.32 | 5.19 |
| CD8 TCM | 0.689 | 1.19 | 0.12 | 6.83 |
| CD4 Tph | 0.159 | 0.39 | 0.07 | 1.32 |
| CD4 TEM_GZMK | < 0.001 | 0.13 | 0.02 | 0.53 |
| CD4 TCM | 0.046 | 0 | 0 | 1.05 |
| CD4 Naive-like | 0.131 | 0 | 0 | 1.4 |
| dnT | 1 | 0 | 0 | 8.64 |

<sup>°</sup>Clonal expansion defined as greater than 5 cells per unique TCRαβ in the peripheral blood dataset and greater than 2 in TIL.
<sup>†</sup> Fisher's Exact test

**Extended Data Table 13.** Association of peripheral blood and TIL single cell clusters with detection in TLS.†.

| Cluster | P value | OR | 95%_CI_Lower | 95%_CI_Upper |
| --- | --- | --- | --- | --- |
| CD4 TEM_GZMB | < 0.001 | 14.12 | 11.35 | 17.72 |
| CD8 TEM_GZMK | < 0.001 | 12.65 | 11.08 | 14.48 |
| CD8 TEM_GZMB | < 0.001 | 10.66 | 9.49 | 12 |
| Treg | < 0.001 | 4.59 | 3.57 | 5.97 |
| NK-T | < 0.001 | 4.58 | 2.08 | 10.85 |
| CD4 TEM_GZMK | < 0.001 | 4.15 | 3.64 | 4.73 |
| MAIT | < 0.001 | 3.09 | 2.33 | 4.11 |
| CD8 TRM | < 0.001 | 2.79 | 1.8 | 4.34 |
| CD4 TCM | 0.012 | 0.89 | 0.81 | 0.98 |
| CD8 TCM | < 0.001 | 0.71 | 0.6 | 0.84 |
| dnT | 0.003 | 0.51 | 0.31 | 0.82 |
| CD4 Tph | < 0.001 | 0.47 | 0.4 | 0.56 |
| gdT | 0.006 | 0.23 | 0.04 | 0.75 |
| CD4 Naive-like | < 0.001 | 0.1 | 0.1 | 0.11 |
| CD4 Naive | < 0.001 | 0.06 | 0.03 | 0.09 |
| CD8 Naive | < 0.001 | 0.03 | 0.01 | 0.05 |

**TIL**
| Cluster | P value | OR | 95%_CI_Lower | 95%_CI_Upper |
| --- | --- | --- | --- | --- |
| CD8 TCM | 0.445 | 1.63 | 0.26 | 7.35 |
| CD4 TCM | 0.505 | 1.52 | 0.34 | 5.48 |
| MAIT | 0.505 | 1.52 | 0.34 | 5.48 |
| CD8 TRM | 0.444 | 1.5 | 0.51 | 3.96 |
| CD4 Naive-like | 0.706 | 1.42 | 0.24 | 6.11 |
| CD4 Tph | 0.812 | 0.76 | 0.22 | 2.15 |
| CD4 TEM_GZMK | 0.567 | 0.72 | 0.28 | 1.66 |
| CD8 TEM_GZMK | 0.23 | 0.7 | 0.39 | 1.22 |
| CD8 TEM_GZMB | 0.472 | 0.6 | 0.15 | 1.84 |
| Treg | 0.003 | 0.22 | 0.07 | 0.66 |
| dnT | 1 | 0 | 0 | 9.1 |

† Fisher's Exact test

**Extended Data Table 14.** Match rate of TLS TCRβ in single cell sequencing of post-treatment peripheral blood and TIL.

| Patient | Total TCRβ, *n* (%) | Top 10% of TCRβ, *n* (%) | Top 1% of TCRβ, *n* (%) | Top 0.1% of TCRβ, *n* (%) |
| --- | --- | --- | --- | --- |
| P02 | 274/11783 (2.3) | 130/1178 (11) | 52/118 (44.1) | 8/12 (67) |
| P03 | 114/10872 (1) | 39/1087 (3.6) | 15/109 (13.8) | 2/11 (18) |
| P07 | 624/11623 (5.4) | 276/1162 (24) | 61/116 (52.6) | 11/12 (92) |
| P08 | 742/33681 (2.2) | 374/3368 (11) | 123/337 (36.5) | 20/34 (59) |
| P12 | 422/7025 (6) | 161/702 (23) | 38/70 (54.3) | 6/7 (86) |
| OT1 | 58/2740 (2.1) | 39/274 (14) | 9/27 (33.3) | 1/3 (33) |
| OT6 | 674/58185 (1.2) | 237/5818 (4.1) | 71/582 (12.2) | 15/58 (26) |
| Total | 2908 / 135909 (2.1) | 1256/13589 (9.2) | 369/1359 (27.2) | 63/137 (46) |

**TIL**
| Patient | Total TCRβ, *n* (%) | Top 10% of TCRβ, *n* (%) | Top 1% of TCRβ, *n* (%) | Top 0.1% of TCRβ, *n* (%) |
| --- | --- | --- | --- | --- |
| OT6 | 196/58185 (0.34) | 135/5818 (2.3) | 76/582 (13.1) | 24/58 (41) |

**Extended Data Table 15.** Summary of antibodies used for immunohistochemistry. Each row indicates a different stain, and columns indicate the target, antigen retrieval buffer used, clone, vendor, product ID and concentration. For dual IHC stains, first and second antibody are indicated.

| Target | Antigen retrieval buffer | First antibody |  |  | Second antibody |  |  |
| --- | --- | --- | --- | --- | --- | --- | --- |
|  |  | Clone | Vendor | Conc. (µg/mL) | Clone | Vendor | Conc. (µg/mL) |
| CD20 | low pH sodium citrate | L26 | Leica | 0.114 |  |  |  |
| CD21/CD3 | low pH sodium citrate | 2G9 | Leica | 4.075 | LN10 | Leica | 0.213 |
| CD8/CD4 | high pH EDTA | 4B11 | Leica | 0.114 | ER204 | Millipore Sigma | 0.096 |
| Ki67/CD20 | high pH EDTA | MM1 | Leica | 0.42 | L26 | Leica | 0.19 |

**Extended Data Table 16.** Summary of antibodies used for FACS.

| Antibody | Clone | Vendor | Dilution |
| --- | --- | --- | --- |
| Live/Dead | Zombie NIR | Biolegend | 1:1000 |
| CD3-FITC | HIT3a | Biolegend | 1:100 |
| CD19-PE/dazzle | SJ25C1 | Biolegend | 1:200 |
| Fc Block | N/A | Biolegend | 1:100 |

